# Gli1 regulates the postnatal acquisition of peripheral nerve architecture

**DOI:** 10.1101/2021.09.14.460314

**Authors:** Brendan Zotter, Or Dagan, Jacob Brady, Hasna Baloui, Jayshree Samanta, James L. Salzer

**Affiliations:** Department of Neuroscience and Physiology, Neuroscience Institute, NYU Langone Medical Center, New York, NY, 10016, USA; Departments of Neuroscience and Clinical Neuroscience, Karolinska Institutet, Stockholm 17177, Sweden; Department of Comparative Biosciences, School of Veterinary Medicine, Stem Cell and Regenerative Medicine Center, University of Wisconsin-Madison, Madison, WI, 53706, USA

**Keywords:** peripheral nerve, Gli1, Schwann cell, myelination, endoneurium, perineurium, desert hedgehog, fate mapping

## Abstract

Peripheral nerves are organized into discrete cellular compartments. Axons, Schwann cells (SCs), and endoneurial fibroblasts (EFs) reside within the endoneurium and are surrounded by the perineurium - a cellular sheath comprised of layers of perineurial glia (PNG). SC secretion of Desert Hedgehog (Dhh) regulates this organization. In Dhh nulls, the perineurium is deficient and the endoneurium is subdivided into small compartments termed minifascicles. Human Dhh mutations cause a peripheral neuropathy with similar defects. Here we examine the role of Gli1, a canonical transcriptional effector of hedgehog signaling, in regulating peripheral nerve organization. We identify PNG, EFs, and pericytes as Gli1-expressing cells by genetic fate mapping. Although expression of Dhh by SCs and Gli1 in target cells is coordinately regulated with myelination, Gli1 expression unexpectedly persists in Dhh null EFs. Thus, Gli1 is expressed in EFs non-canonically i.e., independent of hedgehog signaling. Gli1 and Dhh also have non-redundant activities. In contrast to Dhh nulls, Gli1 nulls have a normal perineurium. Like Dhh nulls, Gli1 nulls form minifascicles, which we show likely arise from EFs. Thus, Dhh and Gli1 are independent signals: Gli1 is dispensable for perineurial development but functions cooperatively with Dhh to drive normal endoneurial development. During development, Gli1 also regulates endoneurial extracellular matrix production, nerve vascular organization, and has modest, non-autonomous effects on SC sorting and myelination of axons. Finally, in adult nerves, induced deletion of Gli1 is sufficient to drive minifascicle formation. Thus, Gli1 regulates the development and is required to maintain the endoneurial architecture of peripheral nerves.

**SIGNIFICANCE STATEMENT:** Peripheral nerves are organized into distinct cellular/ECM compartments: the epineurium, perineurium and endoneurium. This organization, with its associated cellular constituents, are critical for the structural and metabolic support of nerves and their response to injury. Here, we show Gli1 - a transcription factor normally expressed downstream of hedgehog signaling - is required for the proper organization of the endoneurium but not the perineurium. Unexpectedly, Gli1 expression by endoneurial cells is independent of, and functions non-redundantly with, Schwann Cell-derived Desert Hedgehog in regulating peripheral nerve architecture. These results further delineate how peripheral nerves acquire their distinctive organization during normal development and highlight mechanisms that may regulate their reorganization in pathologic settings including peripheral neuropathies and nerve injury.

## INTRODUCTION

Peripheral nerves are organized into distinct compartments (Figure 1A), i.e., the endoneurium, the perineurium, and the epineurium. The organization of these compartments, and their distinct cellular and extracellular matrix (ECM) composition, provides critical structural and metabolic support of nerves and their function in action potential propagation. Cellular components of the endoneurial compartment include axons, Schwann cells (SCs), and endoneurial fibroblasts (EFs), which like SCs are neural crest derived (Joseph et al., 2004). EFs are interspersed between the axon-SC units (Richard et al., 2012) and produce an intervening collagenous ECM. Axon-SC units adopt one of two functional/morphological relationships. Individual SCs ensheath multiple small, unmyelinated axons in separate pockets forming Remak fibers or sort larger axons in a 1:1 relationship, which are then myelinated.

**Figure 1:**
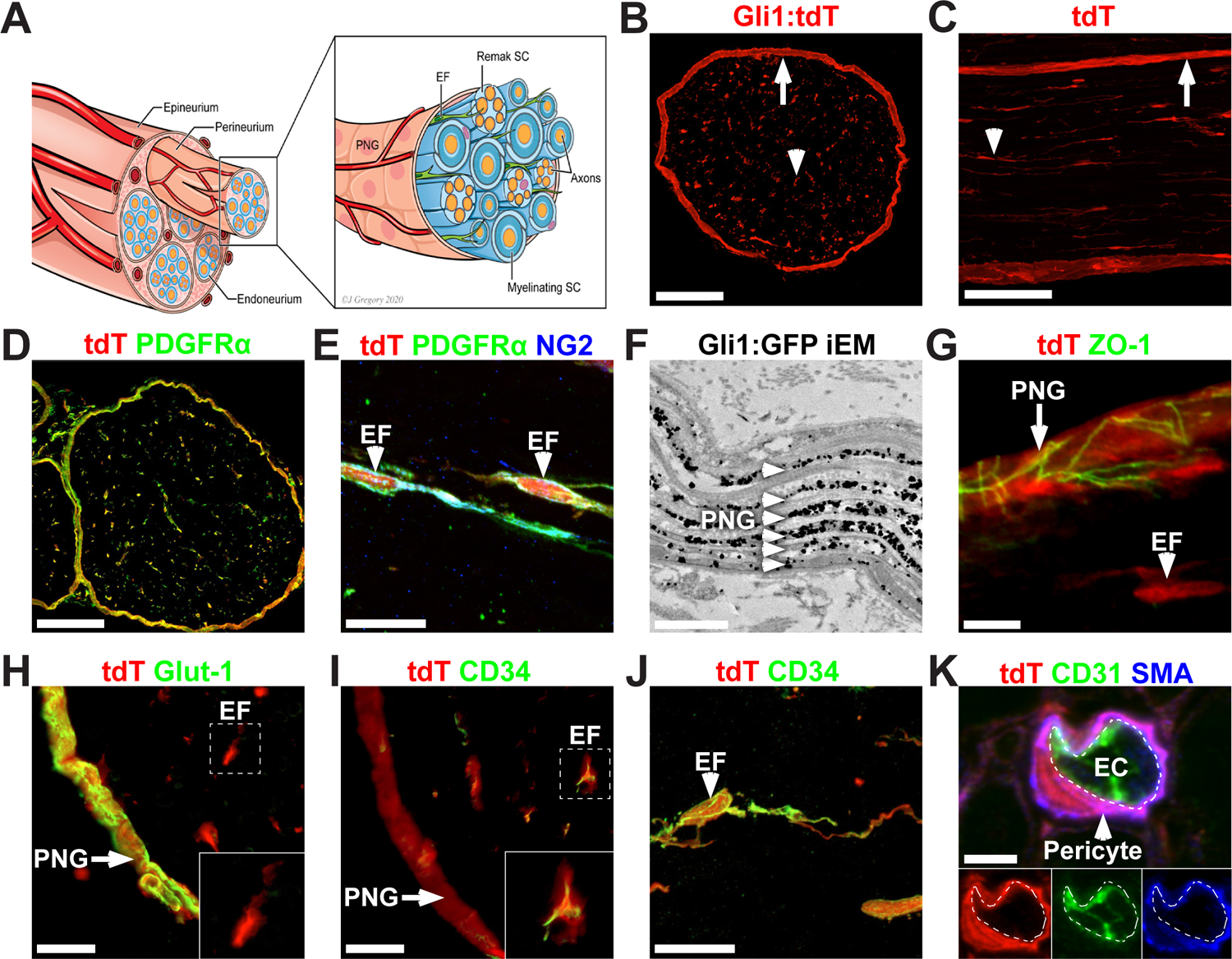
PNG, EF, and pericytes are Gli1-positive PNS cells. (A) Schematic of a peripheral nerve highlighting three cellular compartments: the epineurium, perineurium, and endoneurium. The perineurium is comprised of layers of perineurial glia (PNG), which surround nerve fascicles and delineate the endoneurium, which contains axons (orange) that are either individually myelinated by mSCs (sheath is dark blue) or multiply ensheathed by Remak SCs (light blue), and scattered endoneurial fibroblasts (EF, green). These fascicles are surrounded and bound together by the epineurium. (B) Cross-section and (C) longitudinal section showing fate mapping of the perineurium (arrows) and scattered endoneurial cells (arrowheads) of adult *Gli1^CE/+^;Ai9* (Gli1 het) sciatic nerves. (D) Gli1:tdT fate mapping (red) overlaps almost entirely with PDGFRα (green), a marker of EFs and PNG. (E) EFs labeled with Gli1:tdT (red) co-express NG2 (green), and PDGFRα (blue). (F) Immuno-EM to detect GFP in *Gli1^CE/+^;eGFP* fate mapped nerves labels all PNG layers (arrowheads). The spaces in between labeled cell layers are occupied by collagen fibrils. (G) Fate mapped PNG also express ZO-1 (tight junctions, green) whereas EFs do not (red only). (H) Fate mapped PNG (arrow) express Glut-1 (green), whereas fate mapped EF do not (red only, inset). (I) Fate mapped EF (red) express CD34 (green, inset) whereas fate mapped PNG do not (arrow, red only). (J) High-power view of a fate mapped EF expressing CD-34 (green). (K) A fate mapped (red) endoneurial pericyte is labeled with αSMA (blue) and has no overlap with the endothelial cell (EC) marker CD31 (green). The white dashed line indicates the abluminal surface of the EC. Scale bars: (B-D) 100 µm, (E) 20 µm, (F) 1 µm, (G) 5 µm, (H-I) 10 µm, (J) 20 µm, (K) 3 µm.

This mix of myelinated axons, Remak fibers, EFs, and ECM, are organized into fascicles, which are enclosed by the perineurium. The latter is a cellular sheath and barrier comprised of multiple layers of flattened perineurial glia (PNG), considered to be of neural ectodermal origin (Kucenas et al., 2008; Clark et al., 2014). Finally, PNG-bounded nerve fascicles are further surrounded by the epineurium, consisting of scattered fibroblasts and irregular bundles of collagen fibrils. The epineurium is the outermost sheath that anchors fascicles to each other and to the surrounding mesenchyme (Burkel, 1967). This multilayered fascicular structure provides structural support that limits mechanical strain on axons and SCs while allowing nerves to remain flexible during limb movement (Flores et al., 2000).

The epineurium also contains a plexus of blood vessels, the *vasa nervorum,* that penetrates the perineurium at regular intervals to provide metabolic support to the endoneurium (Bell and Weddell, 1984a). These various components create an immune privileged environment for axon-

SC units (Peltonen et al., 2013). In particular, PNG form tight junctions, creating a perineurial barrier to the invasion of toxins and pathogens (Kristensson and Olsson, 1971). Tight junctions also form between ECs of endoneurial capillaries, constituting the blood nerve barrier (BNB) (Olsson, 1968).

The molecular signals that underlie development of this complex organization, and coordinate assembly of its various components, are incompletely understood. The ensheathment fate of axons, i.e., whether they are myelinated or organize as Remak bundles, is regulated by threshold levels of axonal Neuregulin, together with other signals on the axon and SC basal lamina (Salzer, 2015; Feltri et al., 2016). These signals converge to upregulate a series of SC transcription factors culminating in the expression of Egr2, the master regulator of the SC myelinating phenotype (Topilko et al., 1994). Egr2 controls expression of the structural and metabolic genes necessary for myelination.

Egr2 also directly upregulates expression of Desert Hedgehog (Dhh) by myelinating SCs (mSCs) (Jang et al., 2006). Dhh is known to regulate peripheral nerve organization (Parmantier et al., 1999). Knockouts of Dhh in mice (Parmantier et al., 1999) or mutations of Dhh in patients (Umehara et al., 2000; Baldinotti et al., 2018) result in defective development of the perineurium and epineurium. Loss of Dhh also results in the aberrant formation of minifascicles (MFs), which are small compartments within the endoneurium enclosed by cellular sheaths. MFs are also observed in a number of animal models where SC myelination is impaired, including those with significant reductions of Egr2 and Dhh expression (Darbas et al., 2004; Monk et al., 2011). The cell(s) of origin, the mechanisms of MF formation, and their function remain unknown. Both PNG and EFs have been considered to be candidates to form the MF sheaths (Bunge et al., 1989; Parmantier et al., 1999).

Dhh, along with Sonic Hh (Shh) and Indian Hh (Ihh) comprise the vertebrate Hedgehog (Hh) family - morphogens essential for patterning and development of a variety of tissues (Ingham and McMahon, 2001). All three Hh proteins bind to the Patched (Ptc) receptor on target cells to activate a highly conserved signaling cascade mediated by the co-receptor Smoothened (Smo) and the Gli family of transcription factors (Gli1/2/3) (Briscoe and Thérond, 2013). In the absence of Hh, Ptc tonically inhibits Smo activity and both Gli2 and 3 are constitutively cleaved and function as transcriptional repressors (Gli2R, Gli3R). Hh binding to Ptc relieves its inhibition of Smo, blocking cleavage and resulting in full-length Gli2 and Gli3, which now function as transcriptional activators (Gli2A, Gli3A). Among their transcriptional targets is Gli1 – itself a late transcriptional effector of the Hh pathway (Wilson and Chuang, 2010). Mice that lack Gli1 alone do not exhibit any evident phenotype, likely reflecting compensation by Gli2 (Park et al., 2000).

Gli1 is therefore a candidate transcriptional effector of Dhh activity in peripheral nerves. Moreover, Gli1 expression is considered a sensitive transcriptional readout of Hh pathway activity (Ahn and Joyner, 2004; Dessaud et al., 2008); it is therefore a candidate to both report and mediate Dhh activity in peripheral nerves. However, it is also known that in some settings Gli1 expression is driven via non-canonical pathways, i.e., independent of Hh/Ptc/Smo (Dennler et al., 2007; Po et al., 2017; Hasan et al., 2020).

Here, we used fate mapping to identify EFs, PNG, and pericytes as Gli1-expressing cells in the adult PNS. Gli1 knockout mice recapitulate the MF formation seen in Dhh knockouts, but in contrast have a normal perineurium. We show MFs likely initially assemble from EFs that proliferate and morphologically differentiate in the absence of Gli1. Surprisingly, the EFs that form MFs in Dhh knockouts persistently express Gli1. These latter results strongly suggest Gli1 expression in EFs is non-canonical and that it is required together with canonical Dhh signaling to drive normal development of the endoneurium - as loss of either pathway results in MF assembly. Our findings also implicate Gli1 expression in EFs in ECM production, nerve vascular organization, and modulating the fine structure of axon-SC units. Finally, induced deletion of Gli1 drives MF formation in healthy adult nerves. Together, these results implicate Gli1 in regulating both the development and maintenance of peripheral nerve architecture.

## MATERIALS AND METHODS

### Mouse husbandry

All animal work was conducted under an approved animal protocol and in accordance with the guidelines of the Institutional Animal Care and Use Committee of the New York University School of Medicine. Mice were housed in a temperature- and humidity-controlled vivarium on a 12-hr light-dark cycle with free access to food and water. The following strains were used: *Gli1^CreERT2^* (Jax # 007913), *R26^FSF-CAG-EGFP^* (RCE, Jax # 032037), *R26^FSF-LSL-tdTomato^* (Ai9, Jax # 007909), *Gli1^nLacZ^* (Jax # 008211), *Gli2^nLacZ^* (Jax # 007922), *R26^FSF-FLAG-Gli1^* (Rosa-Gli1, Jax # 013123), *Dhh^-/-^* (Jax # 002784), MPZ-cre (Jax # 017927), *Egr2^flox^* (Generous gift of W.J. Leonard, NHLBI). For lineage tracing of hedgehog-responsive cells, *Gli1^CreERT2^* mice were crossed to either RCE or Ai9 reporter lines. To generate Gli1 null mice, *Gli1^CreERT2/+^* and *Gli1^nLacZ^* mice were crossed to generate *Gli1^CreERT2/nLacZ^* compound heterozygotes. For genetic gain of function studies, *Gli1^CreERT2/+^* were crossed with *R26^FSF-FLAG-Gli1^*.

To generate Egr2 conditional knockouts, *P0-cre* and *Egr2^flox^* were crossed. These animals developed a progressive neuropathy characterized by hindlimb paresis, tremor, and dyscoordinated movement. With supportive care (i.e., nutrient-supplemented gelatin food and presence of littermates for grooming), knockouts survived up to 3 months or longer but were euthanized for humane reasons once they were unable to ambulate using their forelimbs.

### Generation of *Gli1^Flox^* mice

The targeting vector, containing exons 5-9 of the endogenous Gli1 locus flanked by LoxP sites and homology arms. was generated by subcloning from a C57BL/6J BAC library. 10 µg of the targeting vector was linearized and transfected by electroporation into C57BL/6J embryonic stem cells by the NYU Rodent Genetic Engineering Laboratory. After selection with G418 antibiotic, surviving clones were expanded for qPCR analysis to identify recombinant ES clones with single integration. Positive clones were confirmed by southern blot and subsequent sequencing prior to injection into diploid C57BL/6J blastocysts. Resulting chimeric mice were screened by PCR for the presence of the targeted allele and outcrossed to wild-type C57BL/6J mice to confirm germline transmission of the *Gli1^Flox^* allele. The locus was subsequently analyzed by targeted sequencing to confirm fidelity of the modified Gli1 locus. *Gli1^Flox^* mice were crossed to *Gli1^CE^* mice to generate *Gli1^CE/Flox^* compound heterozygotes, i.e., inducible knockouts (iKOs).

### Genotyping

Mice were genotyped by PCR of genomic DNA from tail clips using DreamTaq Green PCR Master Mix (ThermoFisher) and agarose gel electrophoresis. Primer sequences, annealing temperatures (T_a_), and amplicon sizes are listed in the table below. Standard cycle conditions were: 94°C for 3 min, followed by 35 cycles of 94°C for 30 s, T_m_ for 30 s, 72°C for 1 min.

### Genotyping primers

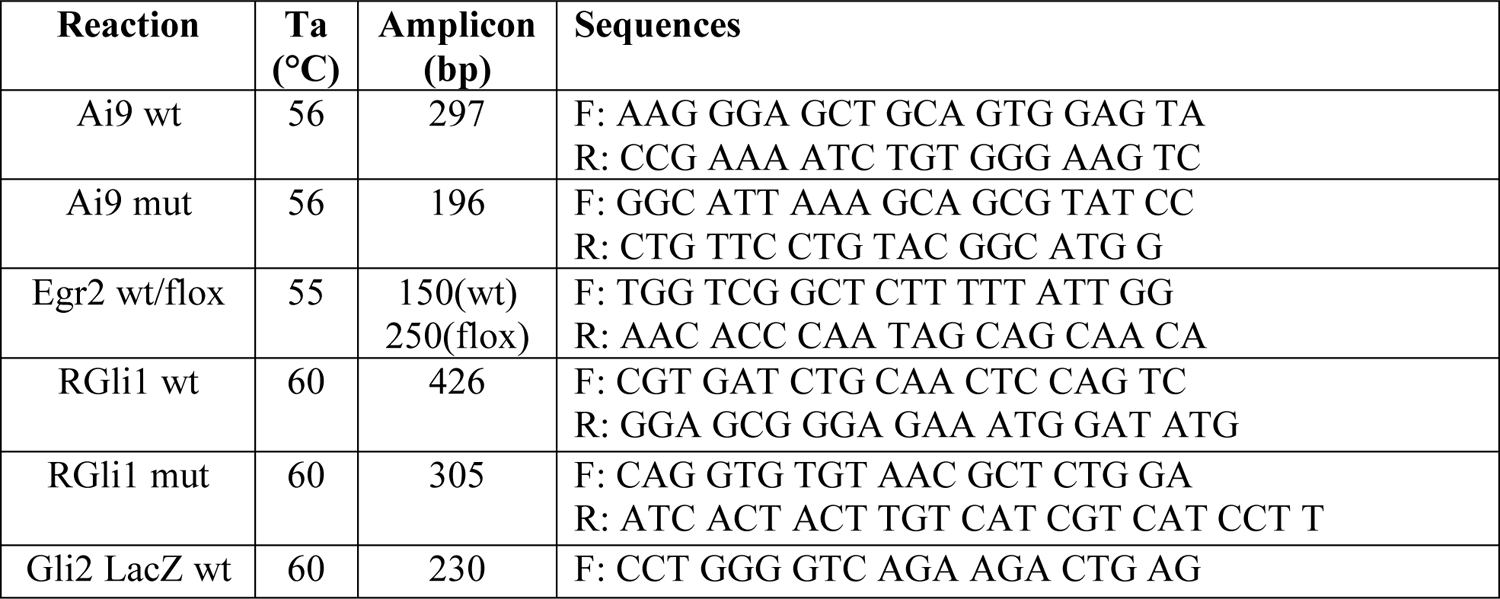
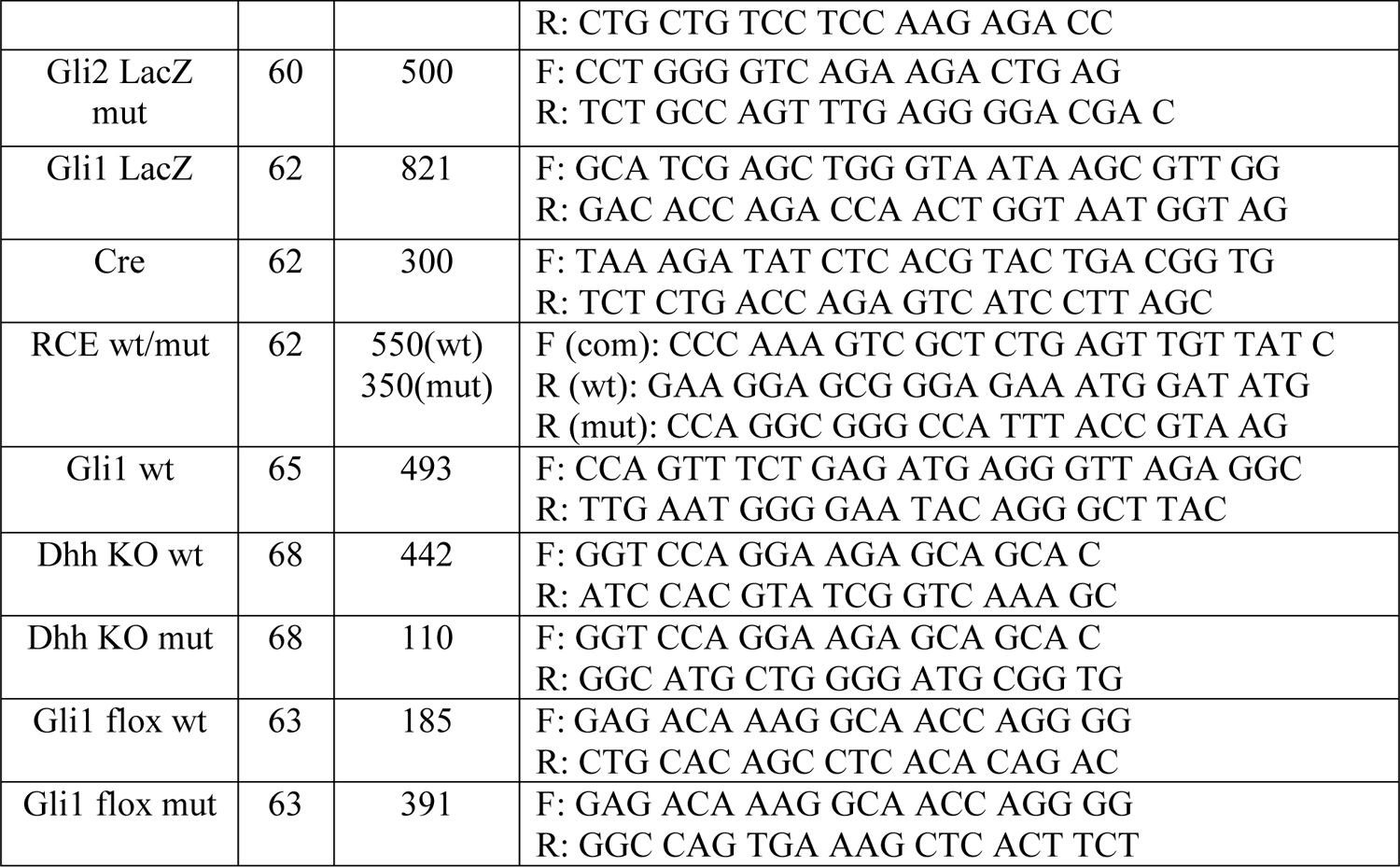

#### Fate mapping

Cre-mediated recombination was induced in adult mice by intraperitoneal (IP) injections of 20 mg/mL Tamoxifen (Sigma) in corn oil (Sigma). 4 injections of 125 mg/kg each were performed on alternating days. For neonates, 5 mg/mL Tamoxifen in corn oil was injected subcutaneously at 150 mg/kg on consecutive days at P0 and P1. For *in utero* fate mapping, pregnant females were given a single dose of 20 mg/mL Tamoxifen at 125 mg/kg by oral gavage at E12 (Danielian et al. 1998). To avoid birthing difficulties due to estrogen receptor blockade, litters were delivered by cesarian section at E18.5 and cross-fostered with other dams with newborn litters (Lizen et al. 2015).

#### EdU Administration

For pulse-labelling, a single injection of 10 mg/mL EdU (Cayman Chemical) in Phosphate-buffered Saline (PBS) at 100 mg/kg was given by IP injection 4 hr prior to tissue collection. Cell proliferation was assessed by EdU incorporation using the Click-iT Plus EdU kit (ThermoFisher) according to the manufacturer’s instructions.

#### Tissue processing

Mice were anesthetized using pentobarbital/phenytoin (Beuthanasia, Merck) and transcardially perfused with PBS, followed by 2% Paraformaldehyde (PFA) in PBS. Sciatic nerves were isolated, pinned to Sylgard 184 (Dow Chemical)-coated plates, and postfixed for 2 hr at 4°C on a rocker. Nerves were cryoprotected in 15% sucrose (ThermoFisher) in PBS overnight, followed by 30% sucrose, and a 1:1 mixture of OCT (Tissue-Tek) to 30% sucrose. Finally, nerves were embedded in 2:1 mixture of OCT to 30% sucrose, flash frozen on liquid N_2_, and sectioned on a Leica CM 3050S cryostat at 12 µm in both longitudinal and cross-sectional planes. Sections were collected on Superfrost Plus slides (ThermoFisher) and dried on a slide warmer at 50°C overnight before storage at −80°C until use.

#### Immunofluorescence

Slides were brought to room temperature and incubated in blocking solution containing 1% Bovine Serum Albumin (BSA, Sigma), 0.25% Triton X-100 (Sigma), and 10% Normal goat serum (Vector Labs) in PBS for 1 hr at room temperature. When goat primary antibodies were included, 10% Normal donkey serum (Jackson Immuno) was used instead. For staining of myelin proteins, slides were permeabilized in Acetone for 20 min at −20°C prior to blocking. After blocking, slides were incubated with primary antibodies diluted in blocking solution overnight at 4°C. Slides were washed in 0.1% Triton X-100 in PBS (PBS-T) for 3 x 5 min, then incubated with appropriate goat or donkey species-specific secondary antibodies (Jackson Immuno) diluted 1:1000 in blocking solution for 2 hr at room temperature. Slides were washed again in PBS-T and mounted with Fluoromount-G (Southern Biotech). To label nuclei, some slides were also incubated in Hoechst 33258 (ThermoFisher) diluted 1:5000 in PBS for 5 min at room temperature prior to mounting. Fluorescent images were obtained with an LSM 800 Scanning Confocal Microscope (Carl Zeiss) using ZEN Blue software (Carl Zeiss). Images were processed using Fiji (NIH) and Photoshop (Adobe). Quantifications were performed on blinded images using counting and tracing tools in Fiji and Photoshop.

### Primary antibodies

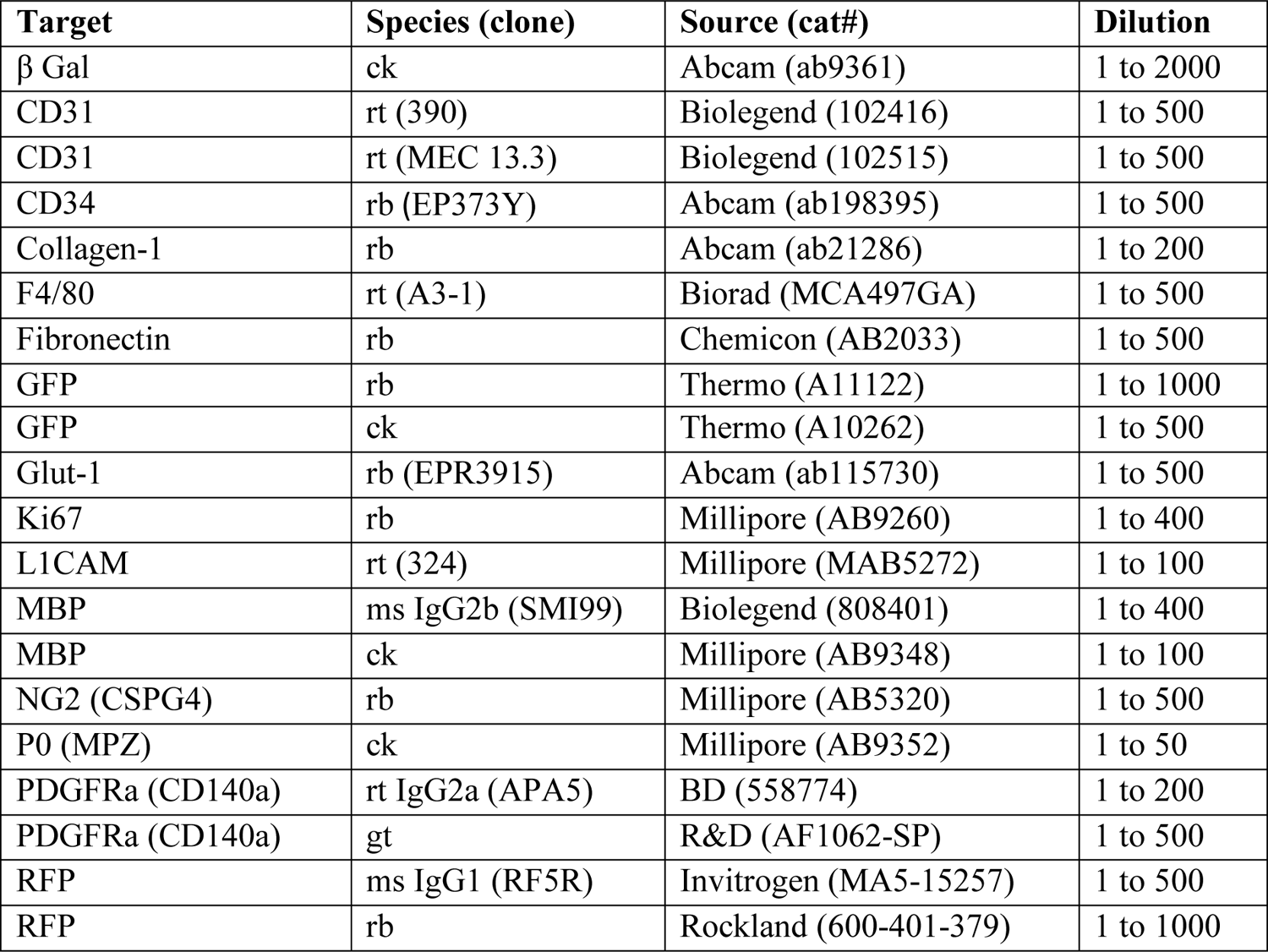
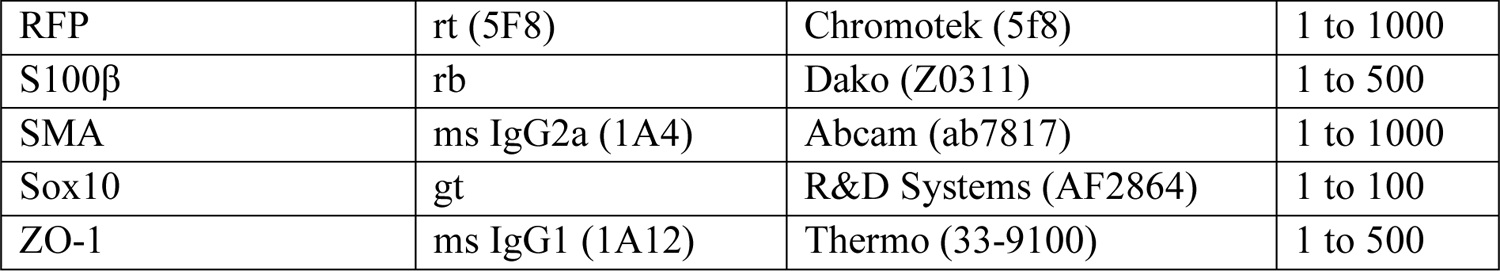

| Target | Species (clone) | Source (cat#) | Dilution |
| --- | --- | --- | --- |
| β Gal | ck | Abcam (ab9361) | 1 to 2000 |
| CD31 | rt (390) | Biolegend (102416) | 1 to 500 |
| CD31 | rt (MEC 13.3) | Biolegend (102515) | 1 to 500 |
| CD34 | rb (EP373Y) | Abcam (ab198395) | 1 to 500 |
| Collagen-1 | rb | Abcam (ab21286) | 1 to 200 |
| F4/80 | rt (A3-1) | Biorad (MCA497GA) | 1 to 500 |
| Fibronectin | rb | Chemicon (AB2033) | 1 to 500 |
| GFP | rb | Thermo (A11122) | 1 to 1000 |
| GFP | ck | Thermo (A10262) | 1 to 500 |
| Glut-1 | rb (EPR3915) | Abcam (ab115730) | 1 to 500 |
| Ki67 | rb | Millipore (AB9260) | 1 to 400 |
| L1CAM | rt (324) | Millipore (MAB5272) | 1 to 100 |
| MBP | ms IgG2b (SMI99) | Biolegend (808401) | 1 to 400 |
| MBP | ck | Millipore (AB9348) | 1 to 100 |
| NG2 (CSPG4) | rb | Millipore (AB5320) | 1 to 500 |
| P0 (MPZ) | ck | Millipore (AB9352) | 1 to 50 |
| PDGFRa (CD140a) | rt IgG2a (APA5) | BD (558774) | 1 to 200 |
| PDGFRa (CD140a) | gt | R&D (AF1062-SP) | 1 to 500 |
| RFP | ms IgG1 (RF5R) | Invitrogen (MA5-15257) | 1 to 500 |
| RFP | rb | Rockland (600-401-379) | 1 to 1000 |
| RFP | rt (5F8) | Chromotek (5f8) | 1 to 1000 |
| S100β | rb | Dako (Z0311) | 1 to 500 |
| SMA | ms IgG2a (1A4) | Abcam (ab7817) | 1 to 1000 |
| Sox10 | gt | R&D Systems (AF2864) | 1 to 100 |
| ZO-1 | ms IgG1 (1A12) | Thermo (33-9100) | 1 to 500 |

#### Masson’s Trichrome Staining

Sciatic nerves were fixed as above, but instead of embedding in OCT, nerves were dehydrated through an ethanol series (70-100%), followed by xylene, and embedded in paraffin wax at 60°C. 5 µm sections were collected on Superfrost Plus slides (ThermoFisher) and dried overnight. Slides were deparaffinized in successive washes of Xylene, 100% ethanol, and 70% ethanol, followed by distilled water. Tissue was stained in Weigert’s iron hematoxylin (Sigma HT1079) for 10 minutes, followed by a 10 minute rinse in distilled water. Next, sections were stained in Biebrich scarlet-acid fuchsin solution (Sigma HT151) for 10 minutes, followed by a 10 minute rinse in distilled water. Stains were developed by incubating in phosphomolybdic-phosphotungstic acid solution (Sigma HT153) for 15 minutes, immediately followed by analine blue (Sigma B8563) for 5 minutes, and 1% acetic acid for 2 minutes. Finally, sections were rapidly dehydrated through an ethanol series followed by xylene and mounted using CC-Mount (Sigma C9368).

#### Barrier Permeability Studies

To test the integrity of the BNB, a solution of 1% Evans Blue (Sigma), 5% BSA (Sigma) in PBS was injected into the tail vein of mice at 10 µL/g body weight. For testing the integrity of the perineurial barrier, 50 µL of the same solution was injected through the fascia over the sciatic nerve in the proximal leg without disturbing the nerve. In both cases, animals were euthanized after 30 min by CO_2_ narcosis and sciatic nerves were rapidly dissected without fixation, embedded in OCT, and flash frozen in liquid N_2_. Nerve cross sections were cut at 12 µm on a Leica CM 3050S cryostat and collected on Superfrost Plus slides. EBA fluorescence was visualized immediately without coverslipping using a Zeiss AxioImager A2 (filter cube: Ex:542-582 nm, Em: 604-644nm).

#### Transmission Electron Microscopy

Mice were anesthetized and underwent trans-cardiac perfusion with 0.1M sodium Cacodylate (EMS) followed by Karnovsky’s fixative (4% PFA, 2% glutaraldehyde, 0.1 M sodium cacodylate, pH 7.4). Nerves were removed and postfixed in Karnovsky’s fixative for at least 72 hr at 4°C with gentle agitation. Samples were then washed in 0.1 M sodium cacodylate, and then post-fixed for 1 hr in 1% osmium tetroxide (EMS) in 0.1 M sodium cacodylate. Nerves were then dehydrated in increasing concentrations of ethanol followed by propylene oxide (PO, EMS). Samples were then infiltrated for 1 h with 1:1 PO:Embed 812 (EMS), and then overnight in 1:2 PO:Embed 812. The following day, samples were transferred to 100% Embed 812 for 2 hr. All incubations were performed at room temperature with gentle agitation. Finally, samples were embedded in 100% Embed 812 and baked at 55°C for 72 hr.

Tissue blocks were sectioned on a Leica UC6 ultramicrotome at a thickness of 1 µm for semithin and 90 nm for ultrathin sectioning. Semithin sections were placed on glass slides and stained with 1% Toluidine blue, 2% Borate in water for 1-2 mins. Ultrathin sections were placed on 2 x 1 mm copper slotted formvar grids (EMS) and counterstained with 3% uranyl acetate in 50% methanol for 20 min followed by lead citrate for 5 min. Grids were viewed on a Talos 120C transmission EM at 120 kV and imaged using a Gatan OneView camera running Digital Micrograph (Gatan). Large tilescans of entire nerve cross sections were acquired using a Zeiss Gemini 300 SEM with Gatan 3View in STEM mode.

#### Calculation of nerve area, axon diameters, and g-ratios

Cross-sectional area of nerves was calculated from Toluidine blue-stained semithin nerve sections. For diameter and g-ratio analysis, at least 10 fields (50 µm x 50 µm) from each biological replicate were quantified using Photoshop (Adobe) to trace individual axons and myelin sheaths, as well as count axons in Remak bundles, fibroblasts, and blood vessels. Axon diameters were derived from the area of the circular traces with a circularity threshold of 0.6. Binning of data was accomplished using custom scripts in Matlab (Mathworks). These scripts will be made available upon request.

### 3View Serial Block-Face Scanning and transmission electron microscopy

For 3View SEM tissue processing, the standard transmission electron microscopy protocol (see above) was used with the following modifications: After Osmium incubation, samples were incubated in 1.5% Potasium Ferrocyanide in 0.1 M CB for 1.5 at RT followed by 1% thiocarbohydrazide for 20 mins at RT. The sample was trimmed and thin sections (70nm) were cut and mounted on slot grids. For STEM imaging, grids were loaded onto the Zeiss grid holder. Images were taken with a beam acceleration of 10.0kV and a working distance of approximately 4.0mm, capturing 8192 pixel by 6144 pixel images with a pixel size of 100 nm to 200 nm. For SBF SEM imaging, the sample block was mounted on an aluminum specimen pin (Gatan, Pleasanton, CA) using silver conductive epoxy (Ted Pella Inc.) to electrically ground tissue block. The specimen was trimmed again and coated with a thin layer of gold/palladium (Denton Vaccum DESK V sputter coater, Denton Bacuum, LLC., NJ, USA). Serial block face imaging was performed using Gatan OnPoint BSE detector in a Zeiss GEMINI 300 VP FESEM equipped with a Gatan 3View automatic microtome unit. The system was set to cut sections with 75 nm thickness, imaged with gas injection setting at 40% (2.9E-03mBar) with Focus Charge Compensation to reduce the charge, and images were recorded after each round of section from the block face using the SEM beam at 1.2 keV with a dwell time of 1.2 µs/pixel. Each frame is 60 x 75 um with pixel size of 3.5 nm. Data acquisition occurred in an automated way using Gatan Digital Micrograph (version 3.31) software. A stack of 150 slices was aligned, assembled using ImageJ, with a volume of 60 x 75 x 15 µm^3^ dimensions was obtained from the tissue block. Segmentation and video were generated by Dragonfly 4.1 (ORS).

#### RNA Isolation and Reverse Transcription

Total RNA was isolated from mouse sciatic nerves using Trizol Reagent (Invitrogen). Flash frozen nerves were pulverized in 1.5 mL Eppendorf tubes over liquid nitrogen using a plastic-tipped electric homogenizer. 1 mL of Trizol was added to each nerve samples and allowed to thaw on ice. Samples were centrifuged at 12,000 G for 12 min at 4°C to pellet lipids and cellular debris. Supernatant was transferred to a fresh tube and processed following a standard Trizol extraction protocol per manufacturer’s instructions with the addition of 15 µg of Glycoblue coprecipitant (Thermofisher) and precipitated overnight at −20°C. 400 ng of total RNA was reverse transcribed in a 40 µL reaction using random hexamers (Promega) and M-MLV Reverse transcriptase (Promega) according to manufacturer’s instructions. All cDNA was then diluted 1:1 in nuclease-free water prior to use in qPCR reactions.

#### Quantitative PCR

Target gene expression was assessed using a BioRad CFX96 thermal cycler in a total reaction volume of 15 µL using PowerUP SYBR Green master mix (ThermoFisher) and 1 µL of cDNA per reaction. Standard qPCR settings were used: 95°C for 10 min followed by 40 cycles of 95°C for 15 s then 60°C for 30 s, followed by melt curve analysis. Ct values for each target gene were internally normalized to the average of at least 2 housekeeping genes (Gln, Hprt1, and CNP1) in each sample and analyzed using the comparative Ct method (Livak and Schmittgen 2001). Specifically, ddCt values for each biological replicate were calculated by subtracting the average control dCT value for that gene from each sample. The resulting normalized ddCT values were log transformed and presented as relative expression values. With this method, control expression is arbitrarily normalized to 1.0.

#### qPCR Primers

For all experiments, primers were designed online using Primer-BLAST (NCBI) and were validated for specificity by Nucleotide-BLAST (NCBI) and the UCSC genome browser (genome.ucsc.edu).

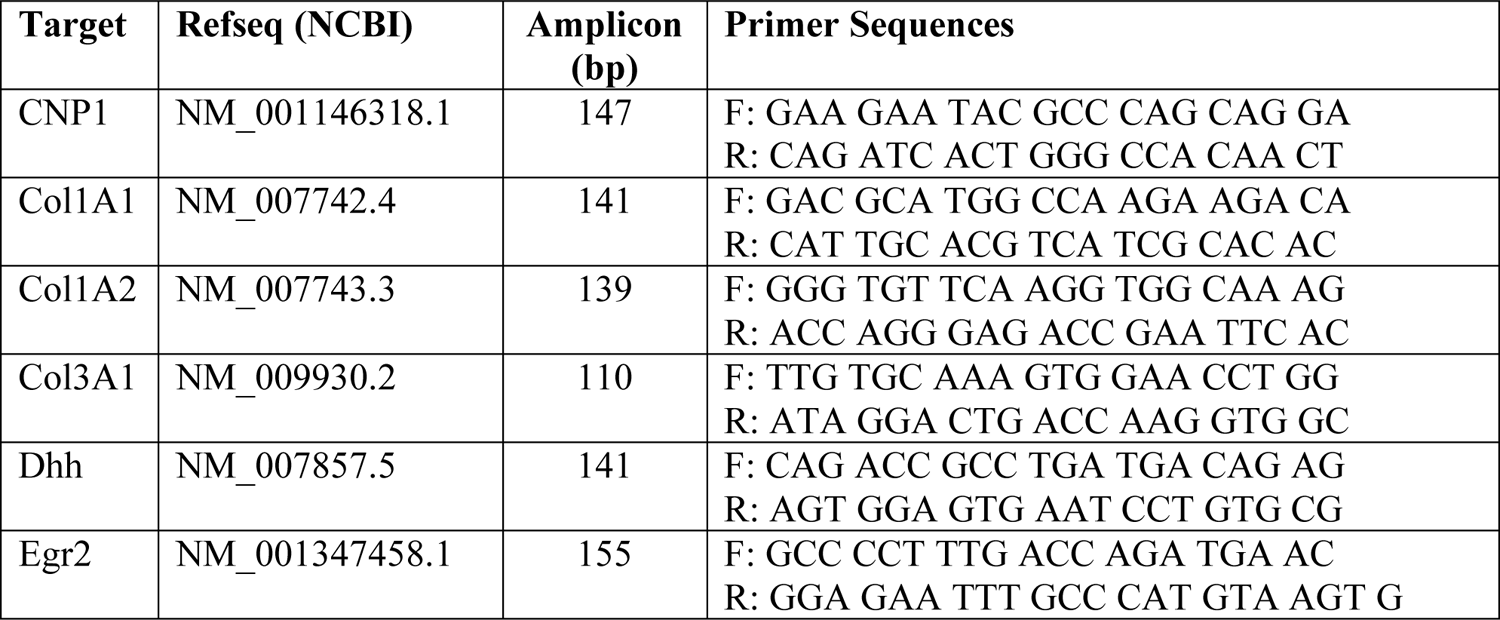
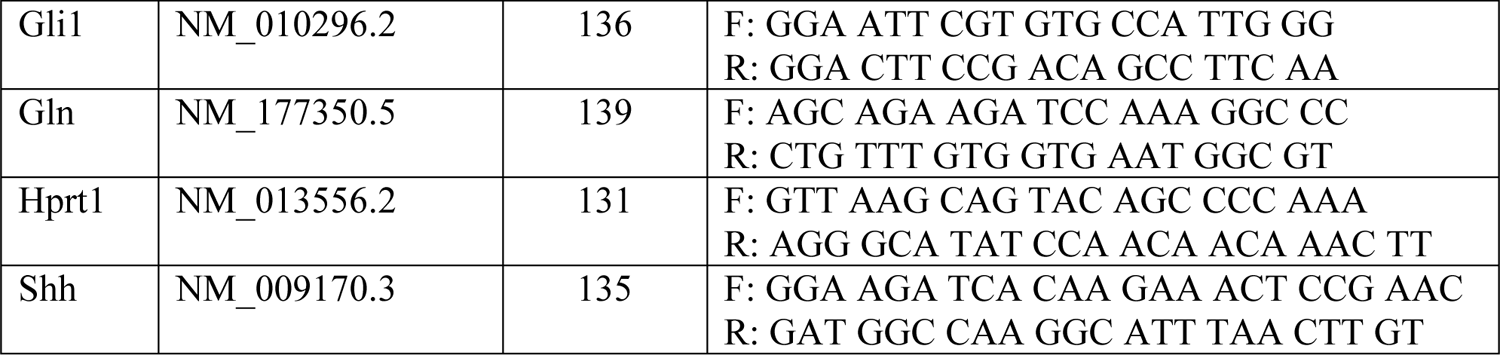

#### Sciatic Nerve Explant Cultures

Gli1 het and null mice were euthanized by CO_2_ narcosis and sciatic nerves were rapidly dissected and placed in cold L-15 media (Sigma). Nerve cultures from each biological replicate were maintained separately. Each nerve was sliced into ∼1 mm segments using a sterile razor blade. segments were transferred to the center of a well in a 6-well tissue culture plate and covered with a minimal (<200 µL) amount of media. A glass coverslip was gently placed on top of the explant to ensure adherence and the well was flooded with 1 mL of EF media consisting of DMEM (Thermo) containing 20% FBS (Gemini). After 5 days, nerve segments and overlying coverslips were removed with sterile forceps and cells that had migrated out from explants were dissociated by addition of 500 µL Trypsin-EDTA (Sigma) to each well. Cells were passaged at a density of ∼5,000 cells/cm^2^ every 7 days for 2 passages to remove any contaminating SCs. Finally, EF cultures were passaged and seeded onto uncoated glass coverslips at a density of ∼5,000 cells/cm^2^ in EF culture media containing either 5 µM GANT61 (MedChemExpress) in DMSO (Sigma) or an equivalent amount of DMSO alone. After 48 hr, cultures were fixed in 2% PFA for 10 minutes and processed for immunofluorescence. For analysis by qPCR, EF cultures were passaged at a density of ∼5,000 cells/cm^2^ in either regular EF or GANT61-containing media in 6-cm plates. After 48 hr, plates were washed in HBSS, incubated with 1 mL Trizol reagent (Invitrogen) at 37°C, and RNA was isolated as above for sciatic nerves with the omission of the initial centrifugation step given the absence of myelin debris. For staining of EF cultures at high and low density, cells were passaged in EF media at either 2,000 or 10,000 cells/cm^2^ on glass coverslips for 48 hr, fixed in 2% PFA for 10 minutes, and processed for immunofluorescence as above.

#### Experimental Design and Statistical Analysis

Mice of both sexes were used in equal ratios whenever possible. All mutants were compared with littermate sibling controls. For all qPCR experiments, 3 biological replicates were prepared for each genotype and all reactions were run in technical triplicate. For all cell counts, proliferation assays, collagen fibril quantification, perineurial thickness, and axon/myelin morphometry, at least 10 individual fields were averaged from each of 3 biological replicates per genotype.

For pairwise comparisons, 2-tailed unpaired t-tests were used with Welch’s correction to account for unequal standard deviations between samples. When multiple comparisons of means were required, Brown-Forsythe and Welch ANOVA tests were used with Dunnet’s T3 multiple comparisons test and a family-wise significance level of 0.05. For assessment of categorical data (such as binned axon diameters), Fisher’s exact test was used to compare numbers of samples failing in each bin with the total number of observations for that sample. For Remak bundle analysis, the number of observations was not sufficient to perform statistical testing by Fisher’s test. All statistical operations and generation of graphs were performed in Prism 8 (GraphPad).

## RESULTS

### Gli1 is expressed in multiple PNS cell types

To identify the full roster of Gli1-expressing cells in the PNS, we fate mapped these cells by crossing *Gli1^CreERT2/+^* mice (Ahn and Joyner, 2004) with the *Rosa26^flox-stop-flox-tdTomato^* (Ai9) reporter (Madisen et al., 2010) to generate *Gli1^CE/+^;Ai9* mice, referred to hereafter as Gli1 hets. Tamoxifen administration in these mice results in permanent labeling of all Gli1-expressing cells and their progeny with cytoplasmic tdTomato (tdT). Fate mapped sciatic nerves from Gli1 hets showed robust tdT labeling of both perineurium (Figure 1B, C, arrows) and cells present throughout the endoneurium (Figure 1B, C, arrowheads).

We found that virtually all fate mapped cells express PDGFR*α*, recently reported to be a broad marker of both PNG and EFs (Richard et al., 2014; Carr et al., 2019) (Figure 1D). Endoneurial fate mapped cells also co-expressed NG2, a known EF marker (Richard et al., 2014) (Figure 1E). Labeling of all cell layers in the perineurium was demonstrated by immuno-EM for a reporter allele driven by Gli1*^CE^* (Figure 1F). Fate mapped perineurial cells also expressed a lattice of ZO-1, a component of the tight junctions that forms between adjacent layers of PNG (Tserentsoodol et al., 1999) (Figure 1G). Finally, we stained nerves for Glut-1/SLC2A1 and CD34, recently confirmed to be selectively enriched in PNG and EFs, respectively in a single cell RNAseq analysis (Gerber et al., 2021). Accordingly, Glut-1 labels fate mapped PNG, but not EFs (Figure 1H) and CD34 labels fate mapped EFs, but not PNG (Figure 1I, J).

There is an additional population of endoneurial Gli1-expressing cells tightly associated with CD31-positive endothelial cells (ECs) that co-expressed *α*SMA (Figure 1K) as well as PDGFR*β* (data not shown). These perivascular cells are likely Gli1-positive pericytes that surround endoneurial capillaries or postcapillary venules (Bell and Weddell, 1984b). All other PNS cell types, including ECs, macrophages, and both mSCs and Remak SCs were Gli1-negative (data not shown). In summary, PNG, EF, and pericytes in peripheral nerves are Gli1-expressing cells based on fate mapping and can be distinguished by cell-type-specific markers.

### Dhh expression by SCs and Gli1 expression in target cells are both regulated by myelination

Dhh expression is largely confined to mSCs (Parmantier et al., 1999). In agreement, transcription of the Dhh gene in SCs is upregulated by the transcription factor Egr2 (Jang et al., 2006), which is required for terminal differentiation of promyelinating SCs into mSCs (Murphy et al., 1996). To determine if Gli1 expression in peripheral nerves, like Dhh, depends on myelination, we generated SC-specific Egr2 conditional knockouts (Egr2 cKO) by crossing *MPZ*-Cre transgenic mice (Feltri et al., 1999) to *Egr2^flox/flox^* mice (Du et al., 2014). These mice were then crossed to *Gli1^nLacZ^* mice, which express nuclear-localized *β*-Galactosidase (*β*Gal) from the endogenous Gli1 locus and therefore provide a readout for active Gli1 expression (Bai et al., 2002). We generated both *Gli1^LacZ/+^;Egr2^flox/+^* (het controls) and *Gli1^nLacZ/+^; MPZ*-Cre; *Egr2^flox/flox^* (cKO) reporter mice. Electron microscopy (EM) analysis showed SCs in the het controls myelinated normally (Figure 2B) whereas SCs in the cKOs were arrested at the promyelinating stage and did not form any myelin (Figure 2D), as expected (Topilko et al., 1994). PDGFR*α* was used to identify PNG and EFs.

**Figure 2:**
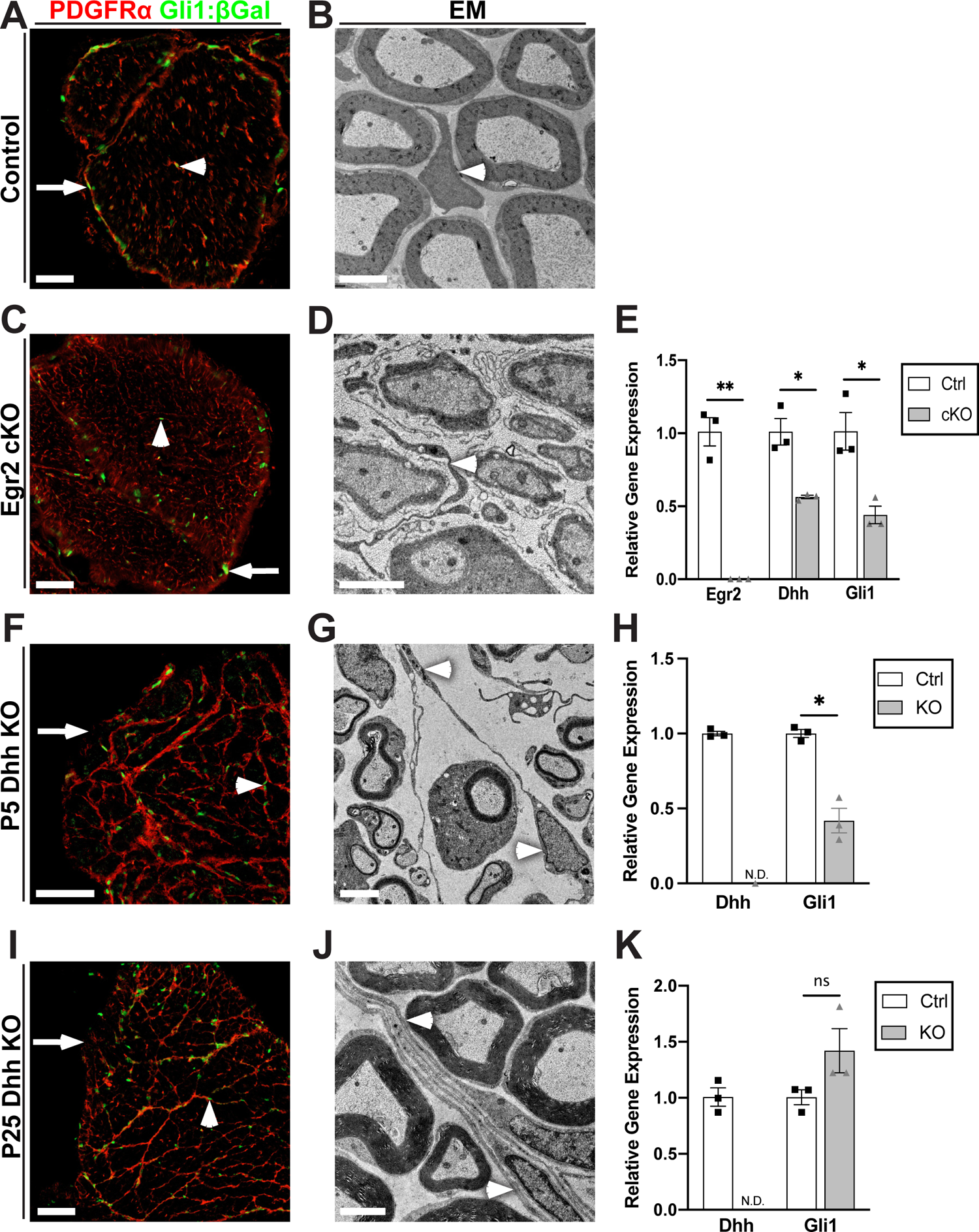
Egr2 regulates Dhh and Gli1 expression but Gli1 expression is independent of Dhh (A) A cross section of *Gli1^LacZ/+^* control nerve stained for *β*Gal (green) and PDGFRα (red) co-labels EFs (arrowhead) and PNG (arrow). (B) EM of an adult control nerve reveals normal morphology of an EF (arrowhead) and of myelinated axons. (C) A cross section from an Egr2 cKO crossed to *Gli1^LacZ/+^* shows *β*Gal/ PDGFRα co-labeling of both EFs (arrowhead) and PNG (arrow). (D) EM shows all axons in the Egr2 cKOs are ensheathed by SCs, which are arrested at the promyelinating stage. By EM, EF morphology is significantly altered with increased process extension (D, arrowhead). (E) qPCR performed on sciatic nerves of P50 controls and Egr2 cKOs shows a >99% reduction in Egr2 mRNA and a ∼50% reduction of both Dhh and Gli1 mRNA. Egr2: Ctrl (1.00 ± 0.97) vs KO (0.002 ± 0.00), p=0.0092; Dhh: Ctrl (1.00 ± 0.91) vs KO (0.56 ± 0.012), p=0.037; Gli1: Ctrl (1.00 ± 0.13) vs KO (0.44 ± 0.06), p=0.03. (F) Cross section from P5 Dhh KO crossed to *Gli1^LacZ/+^* revealed the absence of a perineurium in Dhh KOs (arrow), but the persistence of many Gli1:*β*Gal-positive cells (green, arrowheads) and extensive PDGFRα-positive minifascicles (MFs, red). (G) EM of P5 Dhh KOs reveals normal SC myelination and confirms the reorganization of the endoneurium by nascent MFs (arrowheads). (H) qPCR of P5 controls and Dhh KOs shows Dhh is not detectable and Gli1 expression is reduced to ∼40% of control in Dhh nulls. Dhh: Ctrl (1.00 ± 0.015) vs KO (N.D.); Gli1: Ctrl (1.00 ± 0.027) vs KO (0.42 ± 0.082), p=.013. (I) Cross section from P25 Dhh KO crossed to *Gli1^nLacZ/+^* were stained as above showing similar results as observed at P5. (J) By EM, robust, multilayered MFs are visible throughout the endoneurium. (K) qPCR performed on P25 controls and Dhh KOs shows that Dhh is not detectable, but Gli1 levels are now at or above control levels. Dhh: Ctrl (1.00 ± 0.015) vs KO (N.D.); Gli1: Ctrl (1.00 ± 0.067) vs KO (1.42 ± 0.20), p=.16. Values shown represent mean ± SEM analyzed using unpaired t-test with Welch’s correction based on n=3 biological replicates per genotype; * p<.05, ** p<.01, ns=not significant, N.D.=not detected. Scale bars: (A, C, F, I) 50 µm, (B, D, G, J) 2 µm.

PNG (arrows) and EFs (arrowheads) are *β*Gal-positive in both controls (Figure 2A) and in the Egr2 cKOs (Figure 2C). The density of EFs appeared to be increased with marked process extension compared to controls, but there was not frank MF formation based on staining or EMs (Figures 2C, 2D arrowhead). qPCR of the Egr2 cKOs demonstrated a ∼50% reduction in Dhh and a comparable reduction of Gli1 transcripts (Figure 2E). Together, these results indicate that SC Egr2 expression, which is required for myelination, drives Dhh expression in mSCs and Gli1 expression and phenotypic changes in prospective target cells during postnatal PNS development.

### Gli1 expression in EFs is independent of Dhh

Given the coordinate regulation of Dhh and Gli1 in the Egr2 cKOs, we considered Dhh a potential candidate to drive expression of Gli1 in the PNS. Dhh is also the only member of the Hh family normally expressed in the postnatal PNS (Gerber et al., 2021). In the canonical Hh pathway, Gli1 expression is activated by Gli2A function (Wilson and Chuang, 2010). We therefore first assessed Gli2 expression by crossing *Gli1^CE/+^;Ai9* mice to *Gli2^nLacZ/+^* mice (Bai and Joyner, 2001). This revealed that both endoneurial and perineurial Gli1-positive cells co-expressed Gli2 (Extended data Figure 1), consistent with canonical Hh signaling as a driver of Gli1 expression in these cells.

To examine directly whether Gli1 is downstream of Dhh, we crossed *Gli1^LacZ/+^* reporter mice to *Dhh^-/-^* nulls (Bitgood et al., 1996). In the Dhh nulls, as previously reported (Parmantier et al., 1999), the perineurium is largely deficient (Figure 2F, I, arrows) and there is robust MF formation (arrowheads). Both effects were already evident at P5 based on staining of Dhh null nerves for PDGFR*α* (Figure 2F, I). Elongated cellular processes were observed throughout the endoneurium by EM in P5 Dhh nulls (Figure 2G, arrowheads), with mature MF structures consisting of several cell layers visible by P25 (Figure 2J, arrowheads).

Unexpectedly, Gli1 was persistently expressed in Dhh null sciatic nerves at both P5 (Figure 2F) and P25 (Figure 2I) based on *β*Gal staining. Likewise, by qPCR Gli1 mRNA was expressed in the P5 Dhh nulls, albeit at reduced levels i.e., 56% of controls (Figure 2H). Gli1 levels were normal or even increased in the P25 Dhh nulls compared to controls despite the absence of Dhh (Figure 2K). The amount of Gli1 expression is particularly striking as there is no contribution from Gli1-positive perineurial cells. All *β*Gal-positive cells in the Dhh nulls expressed PDGFRα, indicating that the persistent Gli1 expression remained confined to EFs and was not induced in other cell types, e.g., SCs.

Expression of Gli1 in the Dhh KOs suggests that either other Hh members are upregulated in the Dhh nulls or there is significant non-canonical i.e., Hh-independent induction of Gli1 in EFs. By qPCR, Shh and Ihh transcripts were not detected in P5 Dhh KOs, in Gli1 hets or nulls, or in Egr2 cKOs (data not shown). There may be a small amount of Shh transcripts in the P25 Dhh KOs based on variable detection of an amplicon in PCR cycles 33-38. This varied between biological replicates of the same genotype and even when detected, was much lower than the levels of Dhh in P25 controls, which amplified in cycles 22-23. These results indicate that expression of Gli1, at least in EFs, is largely Hh-independent.

### Gli1 controls endoneurial but not perineurial development

To test the function of Gli1 in the PNS, we generated Gli1 knockout mice by crossing *Gli1^CE/+^;Ai9* mice with *Gli1^LacZ/+^* mice. The resulting Gli1 nulls (*Gli1^CE/LacZ^;Ai9*) were compared to Gli1 hets (*Gli1^CE/+^;Ai9)*; a single copy of *Gli1^CE^* present in both the hets and nulls was used for fate mapping. As previously reported (Park et al., 2000), Gli1 nulls were born at predicted Mendelian ratios and displayed no overt phenotype. qPCR of sciatic nerves confirmed that Gli1 transcript levels in nulls were reduced to less than 5% of the Gli1 hets (data not shown). As Gli1 hets have no evident phenotype and their nerves appeared identical to wild-type nerves by fate mapping and EM (Figure 3A and inset), they were used as controls in all subsequent studies.

**Figure 3:**
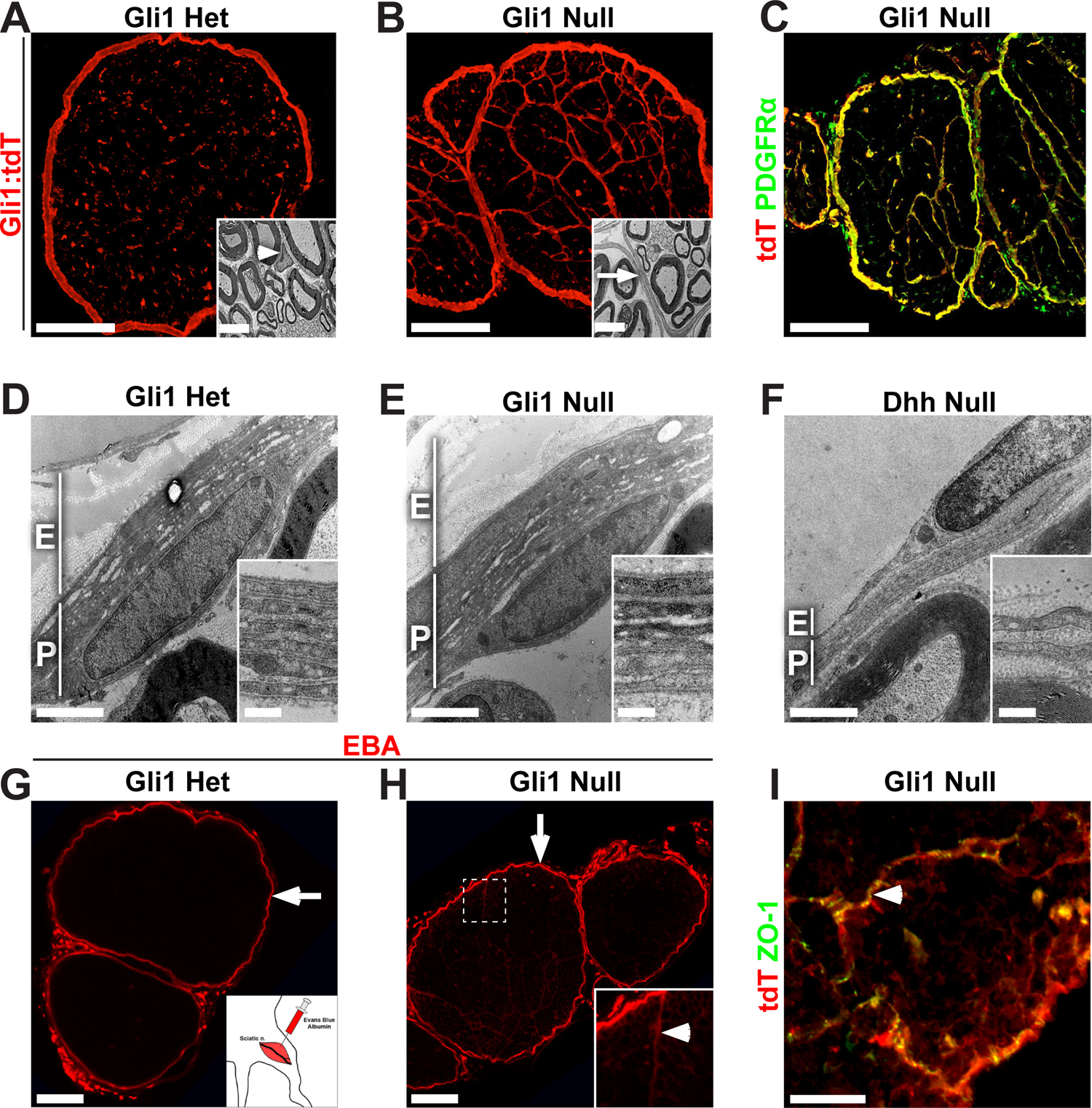
Gli1 controls minifascicle formation but not nerve sheath development (A) Cross section of a fate mapped Gli1 het sciatic nerve reveals numerous tdT-positive endoneurial cells. Inset shows the normal EM morphology of a Gli1 het EF (arrowhead). (B) Cross section of a fate mapped Gli1 null sciatic nerve shows reorganization of the endoneurium into MFs by tdT-positive cells. Inset shows an EM of a MF boundary, comprised of thin, elongated cellular processes. (C) Fate mapping (red) overlaps almost entirely with PDGFRα staining (green), in a Gli1 null. (D, E) EM cross sections through nerve sheath demonstrates comparable thickness of the epineurium (E) and perineurium (P) in Gli1 hets and nulls. Inset shows high power image of concentric layers of PNG. (F) EM cross section of age-matched Dhh null reveals minimal epineurium and a limited perineurium consisting of a few, uncompacted layers. (G) A local injection of Evans Blue-Albumin (EBA, red) was stopped by the perineurial barrier in Gli1 hets (arrow). (H) In Gli1 nulls, most EBA was retained within the perineurium (arrow); small amounts tracked between MF boundaries (inset, arrowhead) which anastomose with the perineurium and allow for dye entry. EBA does not enter the endoneurial compartment. (I) Gli1 null nerves stained for Gli1:tdT and ZO-1 (green) reveals that MFs (arrowhead) have extensive tight junctions, consistent with their intact barrier function. Scale bars: (A-B main) 100 µm, (A-B inset) 5 µm, (C) 100 µm, (D-F main) 2 µm, (D-F inset) 0.5 µm, (G-H) 100 µm, (I) 50 µm.

Nerves from the Gli1 nulls were strikingly organized into MFs, which were formed by thin, closely-associated Gli1 fate mapped processes (Figure 3B and inset). These structures appeared very similar to Dhh null nerves. Both the perineurium and MFs remained PDGFR*α*-positive in the Gli1 nulls (Figure 3C). In contrast to the endoneurium, the epineurium and perineurium in the Gli1 nulls appeared normal in appearance and similar to controls (Figure 3D, E). By EM, the perineurium of Gli1 nulls contained similar numbers of layers as the Gli1 hets i.e., 5.42 ± 0.46 vs. 6.17 ± 0.36, respectively. This contrasts with the Dhh null nerves, which have a minimal epineurium and perineurium (Figure 3F) and, as a result a defective diffusion barrier (Parmantier et al., 1999).

To test if the perineurial diffusion barrier was indeed intact in the Gli1 nulls as suggested by its morphology, we injected Evans Blue Albumin (EBA), a 69 kD fluorescent protein tracer (Wolman et al., 1981), into the local tissue compartment around the nerve. This dye failed to penetrate the perineurium of either the Gli1 hets or nulls (Figure 3G, H, arrows). A very small amount of EBA did track along MF boundaries in the Gli1 nulls, entering at points where MFs anastomose with the overlying perineurium, but did not enter the endoneurial space (Figure 3H, inset). A barrier in the MFs is consistent with the presence of continuous ZO-1 expression between MF-forming cells in the Gli1 nulls (Figure 3I, arrowhead), reflecting the presence of tight junctions. Finally, whereas in Dhh nulls, degenerating/regenerating fibers were noted in aged animals (Sharghi-Namini et al., 2006), none were observed by EM in Gli1 nulls as old as 18 months (data not shown). Thus, proper development of the epi/perineurium and the perineurial barrier requires Dhh but is independent of Gli1 expression.

### Minifascicles arise from progressive remodeling of endoneurial cells

A key question is what cell gives rise to MFs. As MFs share structural features with PNG, including forming tight junctions and a basal lamina, PNG precursors that invade the endoneurial space have been proposed as a source (Parmantier et al., 1999). Alternatively, EFs within the endoneurial compartment may differentiate and acquire properties of the PNG to form MFs. This latter possibility is consistent with the finding that fibroblasts added to myelinating SC/neuron cocultures form perineurial-like structures (Bunge et al., 1989).

To examine the origins of MFs further, we determined when the EF and PNG cell populations arise relative to MF formation. To this end, we carried out *in utero* fate mapping of Gli1 het and null mice by oral gavage of pregnant females at embryonic day 13 (E13) and examined nerves at postnatal day 30 (P30). In hets, this resulted in labeling of both PNG and EFs but not SCs (Figure 4A). Similarly, in Gli1 nulls the MF-forming cells and PNG but not SCs are labeled (Figure 4B). Thus, Gli1-positive precursors to EF and PNG are present early in development and either may potentially contribute to the formation of MFs.

**Figure 4:**
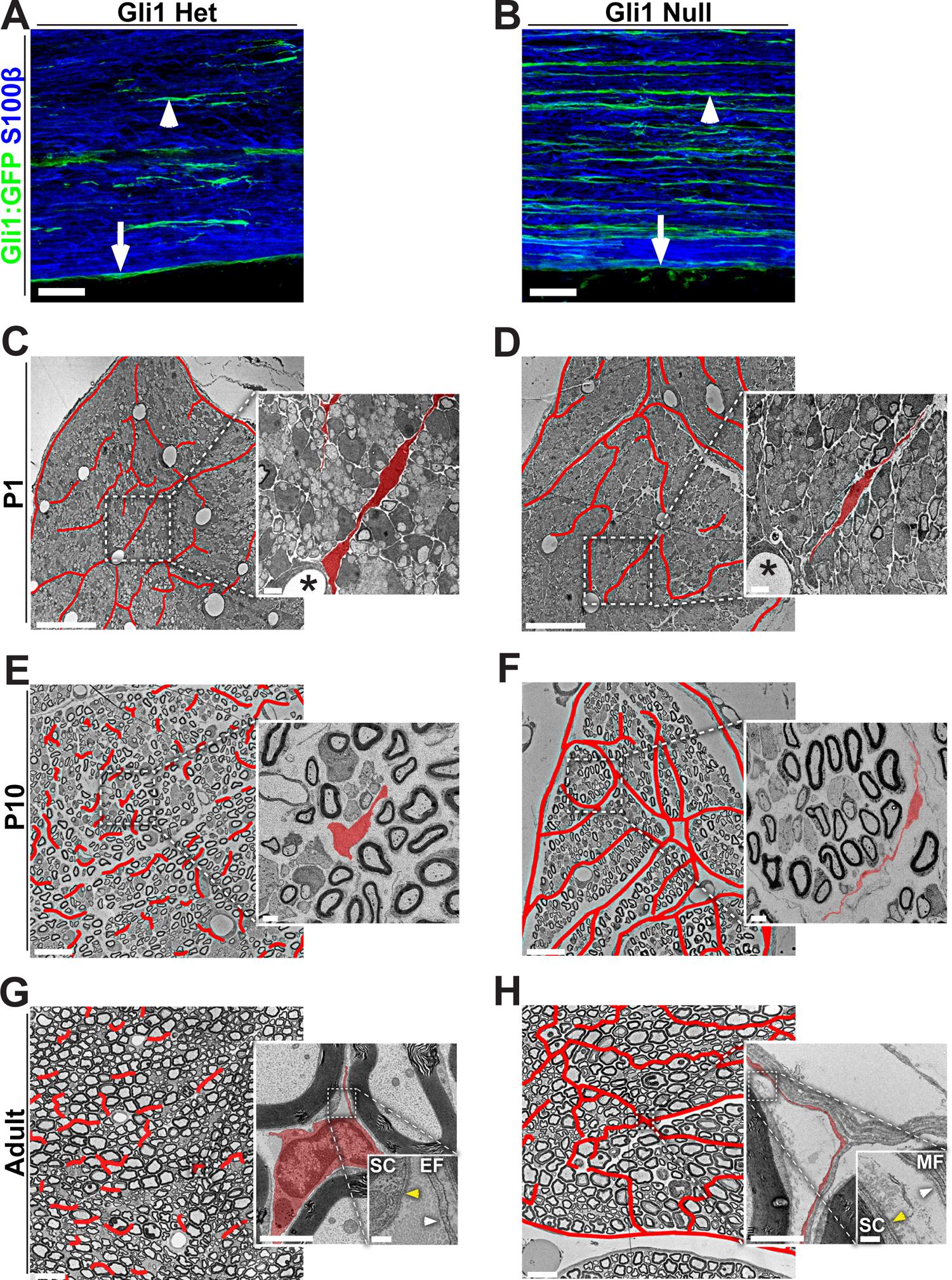
Postnatal development of the endoneurial compartment in Gli1 hets and nulls (A, B) Pregnant Gli1 het and null dams were treated with tamoxifen to fate map embryos of *in utero* at E13. Nerves were harvested at P30, sectioned longitudinally and stained for GFP (green), used as a fate mapping reporter, and for S100β, a marker of SCs (blue). (A) Gli1 hets show fate mapping of EFs (arrowhead) and PNG (arrow), but not SCs. (B) Gli1 nulls show fate mapping of MF-forming cells (arrowhead) and PNG (arrow), but not SCs. (C-H) EM cross sections from Gli1 het and null sciatic nerves from the indicated postnatal times are shown, with endoneurial cells and MFs outlined in red lines. Insets showing fields at higher magnification; individual endoneurial cells are pseudocolored red. (C, D) At P1, little to no myelin is present. Immature EFs in both hets and nulls extend long processes between axon families; many radiate outward from endoneurial blood vessels (insets, stars). (E, F) At P10, robust myelination is observed. EFs in Gli1 hets have retracted their processes and dispersed throughout the endoneurium whereas EFs in Gli1 nulls lengthen processes and begin to wrap around groups of SCs. (G, H) In adult nerves, myelination is complete. In Gli1 hets, EFs are fusiform with short irregular processes whereas in Gli1 nulls the endoneurium is fasciculated by layers of elongated cells. EFs do not form a basal lamina (G inset, white arrow), however a basal lamina is formed by fasciculating cells (H inset, white arrow) and all Schwann cells (G-H insets, yellow arrows). Scale bars: (A-B) 50 µm, (C-H, main) 20 µm, (C-H, insets) 2 µm, (G-H, double insets) 200 nm.

We next characterized when MFs form by analyzing sciatic nerves at different postnatal ages by EM including at P1, P10 and in adults (Figure 4C-H). P1 nerves in both Gli1 hets and nulls contain many EFs, evident as large, flattened cells in the spaces in between developing axon bundles ensheathed by SCs (Figure 4C, D; highlighted in red in insets). At P10 in Gli1 het nerves, these cells begin to remodel, retracting their processes and adopting a more fusiform morphology (Figure 4E). This remodeling continues into adulthood - mature EFs have short processes and lack a basal lamina (4G, white arrow in inset) compared to SCs (4G, yellow arrow in inset); this is consistent with a previous characterization (Richard et al., 2014). In contrast, endoneurial cells in the Gli1 nulls do not retract their processes but rather progressively extend very thin processes around multiple SC/axon units (both mSCs and Remak SCs), evident at P5 (data not shown) and largely complete by P10 (Figure 4F). Their appearance at P10 strongly resembles MFs in adult Gli1 nulls (Figure 4H), including production of a robust basal lamina (see 4H, white arrow in inset). These observations support the notion that MFs arise in a Gli1-dependent fashion, at least in part via remodeling of immature EFs between P1-P10.

### EF but not PNG proliferation is increased in Gli1 nulls

Formation of MFs is associated with an increase in the numbers of its constituent cells. We therefore counted Gli1-positive EF and PNG, indicated by positive *β*Gal immunostaining in *Gli1^LacZ/+^* hets and *Gli1^CE/LacZ^* nulls; Sox10 served as a marker of SCs (Figure 5A, B). There were roughly twice as many *β*Gal-positive, Sox10-negative cells (i.e., EF/PNG) in Gli1 nulls vs hets starting at P5 - an increase that persisted into adulthood (Figure 5C). To determine if this increase resulted from increased cell proliferation in the Gli1 nulls, we pulse labeled with EdU at various developmental ages and counted EdU-positive cells in the endoneurium and perineurium. This revealed an increase in EdU incorporation in *β*Gal-positive, Sox10-negative cells in the Gli1 nulls from P1 to P10. The increase in proliferation was exclusively in the endoneurium, not the perineurium (Figure 5D) supporting the notion that MF formation originates from the proliferation and reorganization of EFs. In further support, there were fewer “free” EFs (i.e., single cells within the endoneurium) in adult Gli1 null nerves, suggesting their disappearance from the endoneurium reflects their accumulation into MFs (Figure 5E-G). By P15 only a few EdU+ cells were detected in either genotype and none were observed in adults (data not shown) consistent with MF formation being largely established by P10 (see Figure 4) and that once formed, these structures are stable.

**Figure 5:**
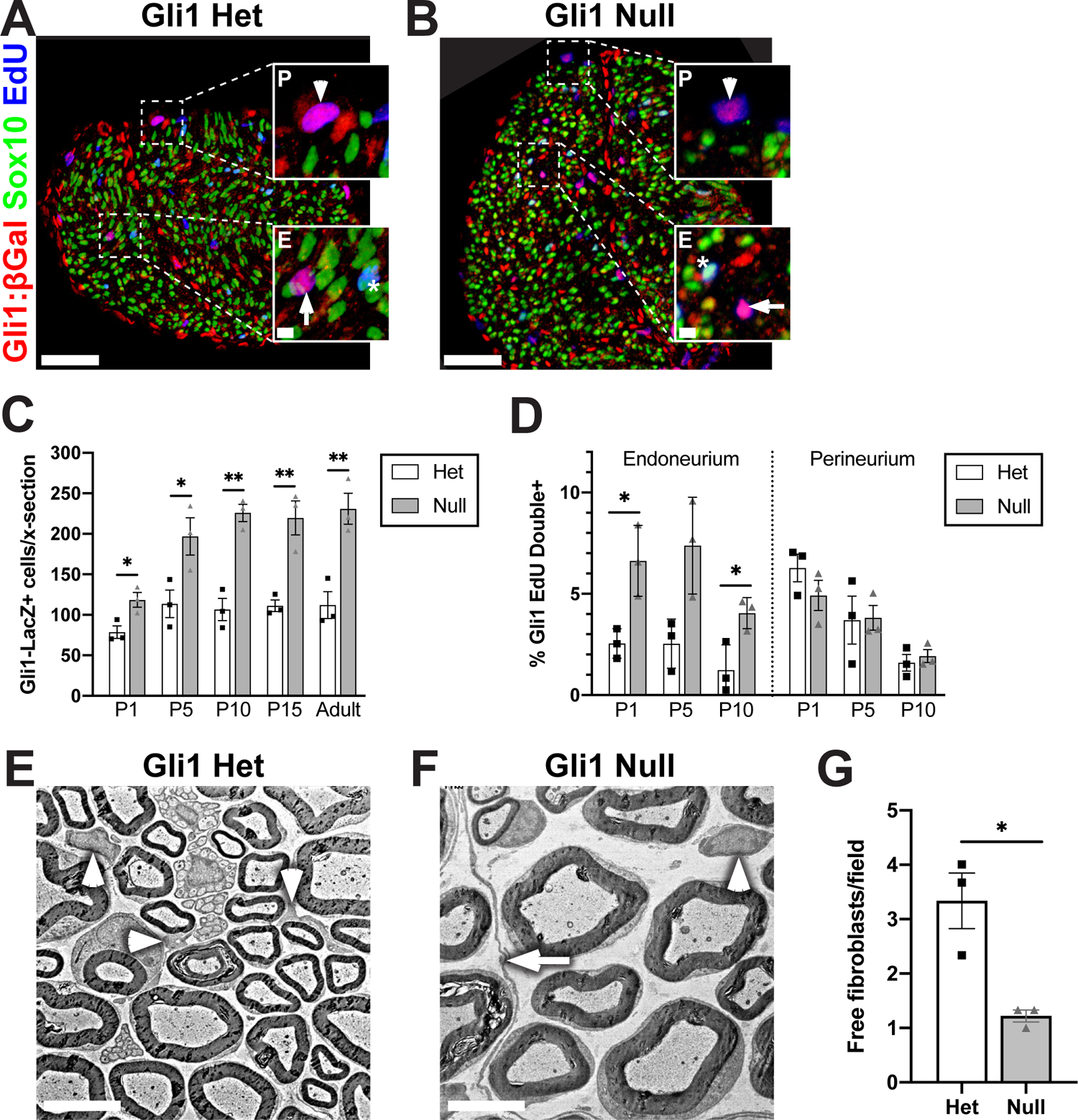
Loss of Gli1 increases endoneurial fibroblast proliferation *in vivo* (A, B) Cross sections of P5 Gli1^LacZ/+^ het and Gli1^CE/LacZ^ null sciatic nerves were stained for *β*Gal (red), Sox10 (SCs, green), and EdU (blue) to mark proliferating cells. <u>Insets:</u> Dividing Gli1 cells (magenta) are found in the endoneurium (E, arrows) and perineurium (P, arrowheads). Dividing SCs are cyan (stars). (C) Gli1-*β*Gal-positive cells were increased in Gli1 nulls vs hets at all postnatal timepoints. P1: Het (78.7 ± 7.7) vs Null (118.4 ± 9.2), p= 0.029; P5: Het (113.7 ± 17.0) vs Null (196.9 ± 23.0), p=0.044; P10: Het (106.6 ± 13.7) vs Null (225.8 ± 10.7), p=0.0024; P15: Het (111.2 ± 7.1) vs Null (219.7 ± 21.1), p=0.0082; Adult: Het (112.0 ± 16.6) vs Null (231.0 ± 19.1), p=0.0093. (D) Proliferation of Gli1-*β*Gal-positive cells as measured by EdU incorporation is increased in endoneurial but not perineurial cells in Gli1 nulls vs. Gli1 hets at the times indicated. Endoneurium: P1: Het (2.5 ± 0.42) vs Null (6.6 ± 1.0), p=0.041; P5: Het (2.5 ± 0.70) vs Null (7.4 ± 1.4), p=0.053; P10: Het (1.2 ± 0.72) vs Null (4.0 ± 0.44), p=0.038; Perineurium: P1: Het (6.3 ± 0.68) vs Null (4.9 ± 0.74), p=0.25; P5: Het (3.7 ± 1.2) vs Null (3.8 ± 0.61), p=0.94; P10: Het (1.6 ± 0.41) vs Null (1.9 ± 0.32), p=0.56. (E, F) EM of adult Gli1 het and null sciatic nerves highlighting “free” EFs (arrowheads), i.e., isolated from any other EF/PNG cell processes. MFs are visible in Gli1 nulls (arrow). (G) Free EFs are reduced in the Gli1 nulls (1.22 ± 0.11) vs Gli1 het (3.33 ± 0.51) per field, p=0.0479. Values shown represent mean ± SEM analyzed using unpaired t-test with Welch’s correction based on n=3 biological replicates per genotype; * p<.05, ** p<.01. Scale bars: (A-B, main) 50 µm, (A-B, insets) 5 µm, (E-F) 5 µm.

To confirm that Gli1 expression regulates EF proliferation in a cell-autonomous fashion, we characterized fate mapped cells derived from sciatic nerve explants of Gli1 hets and nulls. Spindle-shaped tdT-positive cells rapidly migrated out of the ends of het and null sciatic nerve segments explanted into culture. These cells were passaged in high-serum media without additional growth factors, conditions which enhance survival of EF/PNG and eliminates SC contamination (Ochoa and Spinel, 2010). In such cultures, the tdT cells gradually adopted a flattened morphology, and expressed fibronectin and ZO-1, further establishing them as EFs and/or PNG (Peltonen et al., 1987) (data not shown). We then counted the number of tdT-positive cells 48 hr after plating and also stained for Ki67 to mark mitotic cells (Figure 6A-C). We found a significant increase in cell density (Figure 6D) and proliferation (Figure 6E) in knockout cultures compared to controls. Further, addition of 5 µM GANT-61, a pharmacological inhibitor of Gli1 and Gli2 activity (Lauth et al., 2007), increased cell density (Figure 6D) and cell proliferation (Figure 6E) in control cultures to the same levels as the Gli1 nulls. The efficacy of GANT-61 treatment was corroborated by the significant reduction in Gli1 transcript levels (Figure 6F); there was also a ∼25% reduction in Gli2 transcript levels, which did not quite reach significance (data not shown).

**Figure 6:**
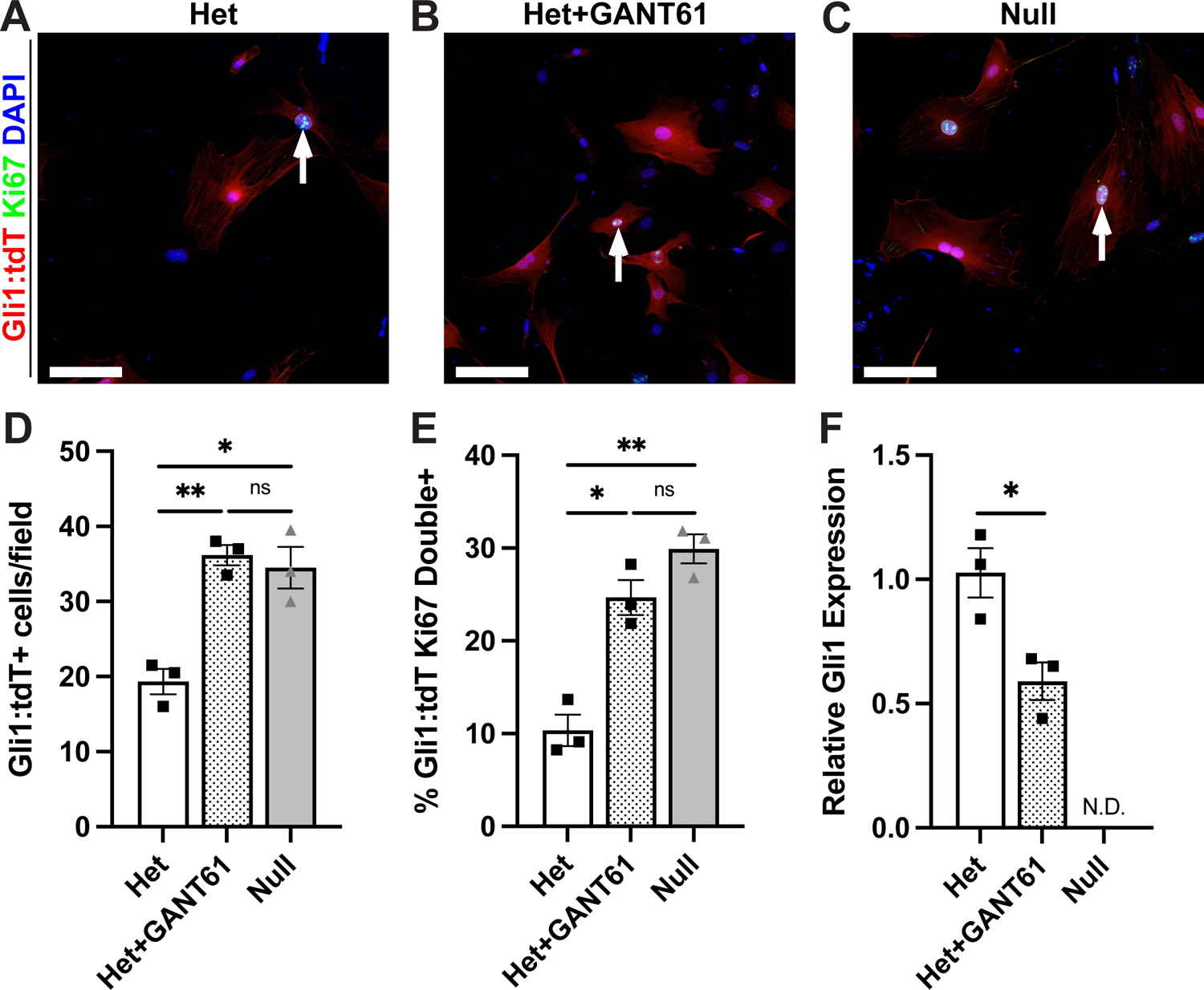
Gli1 loss drives EF/PNG proliferation *in vitro* (A-C) Gli1-fate mapped cells, established from sciatic nerve explants of Gli1 het and nulls, were cultured for 48 hr in either control media or media containing 5 µM GANT61, an inhibitor of Gli1 and Gli2 activity. Cells were fixed and stained with antibodies to the tdT reporter (red), Ki67 (green), and Hoechst (nuclei, blue). Dividing EF/PNG are red with cyan nuclei (arrows). (D) The total number of tdT-positive cells per field was increased in Gli1 nulls (34.5 ± 2.75) and in Gli1 hets + GANT61 (36.17 ± 1.36) compared to Gli1 het controls (19.33 ± 1.69); Het vs Het+GANT61 p=0.0038, Het vs Null p=0.043, Het+GANT61 vs Null p=0.92. (E) The proliferation rate, as assessed by Ki67 staining, was increased in Gli1 nulls (29.91 ± 1.57) and Gli1 hets + GANT61 (24.68 ± 1.88) vs Gli1 het controls (10.35 ± 1.70) cultures; Het vs Het+GANT61 p=0.012, Het vs Null p=0.0027, Het+GANT61 vs Null p=0.23. Values shown represent mean ± SEM analyzed using Welch’s ANOVA test with Dunnett’s T3 multiple comparisons test based on n=10 fields across multiple coverslips per condition and n=3 biological replicates per genotype (prepared from separate nerves); * p<.05, ** p<.01, ns=not significant. (F) qPCR performed on cultured cells confirms suppression of Gli1 by GANT61 treatment. Het (1.02 ± 0.16) vs Het+GANT61 (0.66 ± 0.11), p=.029; Gli1 null (N.D.). Values shown represent mean ± SEM analyzed using paired t-test based on n=3 biological replicates per genotype; * p<.05, N.D.=not detected. Scale bars: (A-C) 100 µm.

### Cell marker analysis implicates EFs in MF assembly

To address further which cells give rise to MFs, we stained perinatal and adult sciatic nerves of Gli1 (Figure 7B, E, H, K) and Dhh (Figure 7C, F, I, L) nulls for CD34 and Glut-1, markers enriched in EFs and PNG in wild type nerves, respectively (see Figure 1H-J). Nerves were also stained for PDGFR*α* to demarcate cells corresponding to the Gli1-fate mapped population (see Figures 1D, 3C). This analysis showed that MFs forming in early postnatal (P5) Dhh and Gli1 nulls (panels A-F) were almost exclusively comprised of CD34-positive, Glut-1-negative cells, providing further evidence that forming MFs arise from EFs and not PNG.

**Figure 7:**
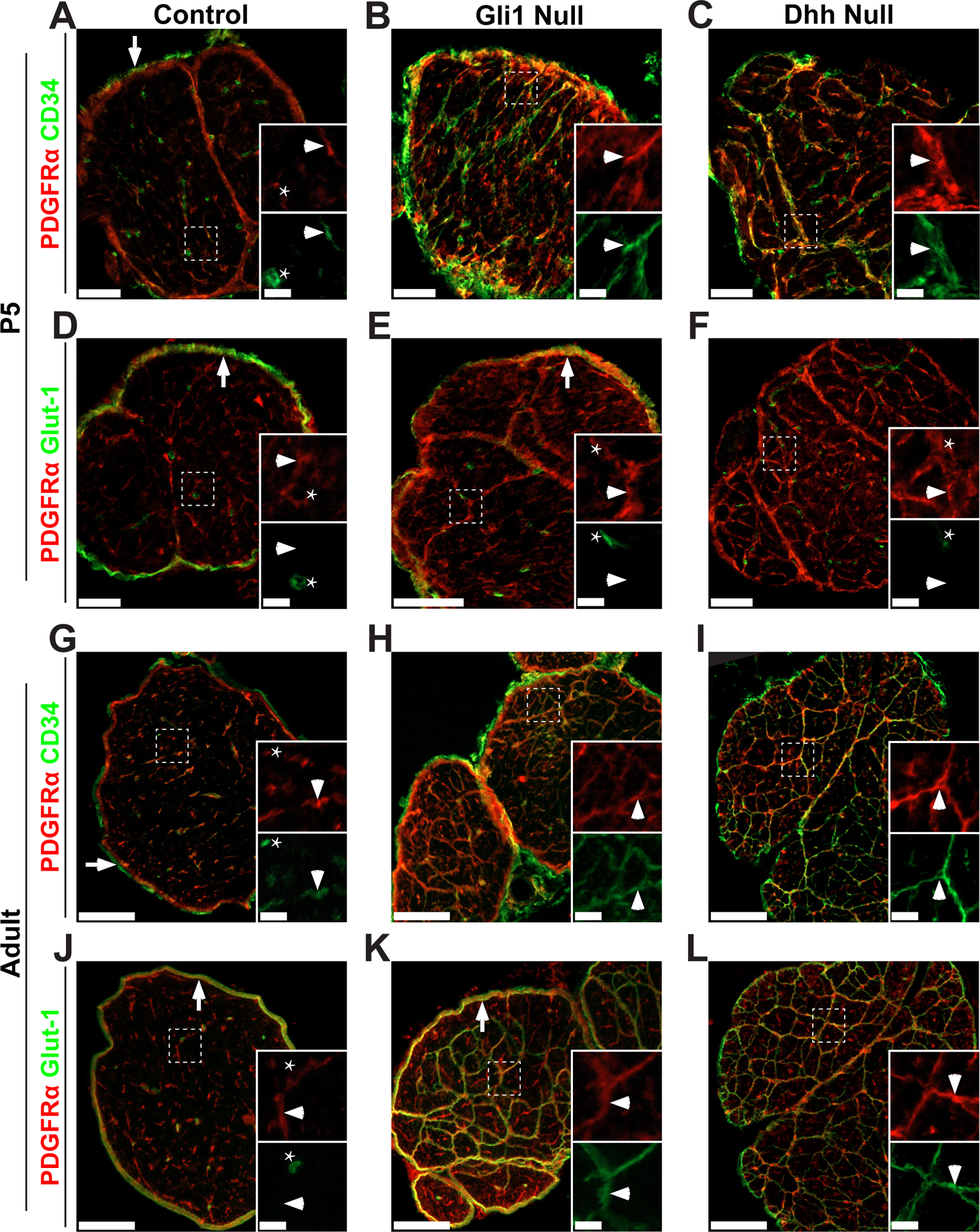
Minifascicles in Gli1 and Dhh nulls express only an EF marker at P5 but express both EF and PNG markers in adult nerves Cross sections of P5 and adult control (A, D, G, J), Gli1 null (B, E, H, K) and Dhh null (C, F, I, L) nerves show staining for PDGFRα (red) and either CD34 (A-C, G-I) or Glut-1 (D-F, J-L), (both green). Insets are higher magnifications of the boxed area showing separated channels. (A) P5 control nerve inset shows labeling of an EF (arrowhead) and an EC, which expresses CD34 but not PDGFRα (star). Epineurial fibroblasts also express CD34 (arrow). In P5 Gli1 null (B) and Dhh null (C) nerves, nascent MFs are extensively labeled by CD34 (insets, arrowhead). (D) P5 control nerves shows Glut-1 is expressed by PNG (arrow) but not EFs (inset, arrowhead). ECs also express Glut-1 but not PDGFRα (inset, star). In P5 Gli1 null (E) and Dhh null (F) nerves, nascent MFs are not labeled by Glut-1 (insets, arrowhead). The Glut-1 positive perineurium present in Gli1 nulls (E, arrowhead) is absent from in Dhh nulls. (G) Adult control nerve stained shows labeling of EFs (inset, arrowhead), ECs (inset, star), and epineurial fibroblasts (arrow). In both adult Gli1 null (H) and Dhh null (I) nerves, mature MFs continue to express CD34 (insets, arrowhead). (J) Adult control nerve for Glut-1 shows labeling of PNG (arrow) and ECs (inset, star), but not EFs (inset, arrowhead). In adult Gli1 null (K) and Dhh null (L) nerves, mature MFs express high levels of Glut-1. Gli1 nulls also have a robust Glut1-positive perineurium that surrounds the MFs (K, arrow) which is largely absent in the Dhh nulls. Scale bars: (A-F, main) 50 µm, (A-F, insets) 4 µm, (G-L, main) 100 µm, (G-L, insets) 20 µm.

In contrast, in the adult Dhh and Gli1 nulls (panels G-L), cells that comprise MFs express both CD34 and Glut-1. This may reflect that these MFs contain a mixture of cells that express either CD34 or Glut-1 or, alternatively that a single population of CD34-positive cells upregulates Glut-1 as MFs mature. We were unable to double stain nerves to determine if Glut-1 and CD34 are co-expressed or are in separate cell populations as, based on our testing, the most reliable commercially-available antibodies to these markers are both raised in rabbit. However, the widespread expression of both markers in the MFs of both Gli1 and Dhh nulls (Figure 7) and the relatively uniform cytoarchitecture of these structures by EM (Figure 4H) suggests adult MFs are comprised of a single population of cells that co-express both markers.

We next examined if expression of these markers, particularly Glut-1, is upregulated in adult MFs potentially as the result of increased cell density and cell contact characteristic of these adult structures. We therefore analyzed cultures of fate mapped cells from Gli1 het and null sciatic nerves grown at low and high cell density (Extended data Figure 2). We found that CD34 expression is higher when these cells are grown at lower density, whereas Glut-1 expression, visible along the membranes of cells in direct contact, is only expressed when cells are grown at high density. The staining data, taken together with the EM and proliferation studies described above, support EFs as the likely cell of origin for MFs in both Gli1 and Dhh nulls. They also suggest these structures eventually acquire perineurial features including Glut-1 expression, possibly as a result of increased cell-cell contact.

### Gli1 regulates endoneurial architecture

We next examined whether there were other changes in nerve architecture in the Gli1 nulls, in addition to MFs, focusing on the vasculature, the ECM, and axon-SC units. 3D reconstruction of nerves from Gli1 nulls by serial block-face scanning EM (SBF-EM) showed that most endoneurial blood vessels are embedded within MFs (Extended data video 1). Staining cross sections of nerves for CD31 to label ECs confirmed that Gli1-positive cells were closely associated with blood vessels in both Gli1 hets and nulls (Figure 8A, B, insets). We also saw a significant increase in the overall number of blood vessels in Gli1 nulls compared to het controls (Figure 8C). The increased blood vessel density and the close spatial relationship of Gli1-positive cells with blood vessels was evident at all timepoints starting at P1 (Figure 8D, E), suggesting an increase in developmental vascularization rather than neovascularization occurring in adults. The BNB was grossly intact as intravenously-injected EBA was confined to blood vessels and the narrow space within fascicle walls, never entering the interior of the fascicles in Gli1 nulls (Figure 8F, G). In both genotypes, EBA leaked out from epineurial capillaries, which are known to be fenestrated and lack tight junctions (Olsson, 1990). We also monitored macrophage numbers by crossing Gli1 hets and nulls to CX3CR1^EGFP/+^ reporter mice (Jung et al., 2000) and saw no significant difference in macrophage numbers between genotypes (Figure 8H-J). Together, this data supports a role of Gli1 expression in endoneurial cells non-autonomously regulating development of the nerve vasculature, but not BNB integrity.

**Figure 8:**
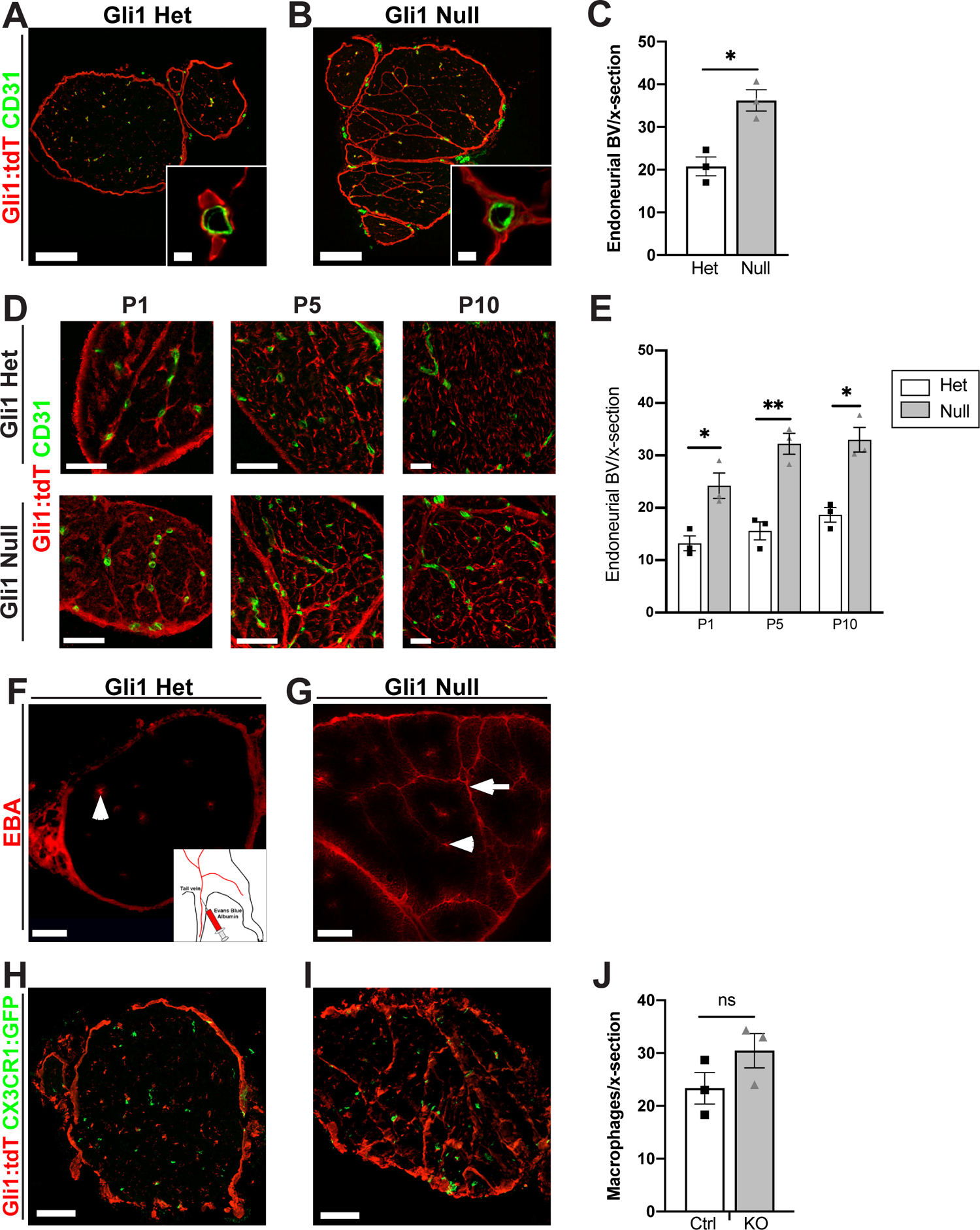
Gli1 expression controls peripheral nerve vascularization (A, B) Sciatic nerve sections from adult fate mapped Gli1 hets and nulls (tdT, red) were stained for the EC marker CD31 (green). Vessels are surrounded by Gli1-positive cells in both hets and nulls. (C) Density of CD31+ vessels is significantly increased in adult Gli1 nulls (36.22 ± 2.50) vs hets (20.78 ± 2.21), p=0.010. (D) Nerve cross sections from Gli1 het or null mice at the indicated postnatal timepoints were stained for tdT and CD31. Blood vessels are closely associated with tdT-positive cells starting at P1 in both genotypes; MFs surround endoneurial vessels in Gli1 nulls. (E) Quantification shows vasculature is significantly increased in Gli1 nulls starting at P1 and persisting into adulthood. P1: Het (13.22 ± 1.42) vs Null (24.22 ± 2.41), p=0.025; P5: Het (15.63 ± 1.71) vs Null (32.22 ± 2.00), p=0.0035; P10: Het (18.67 ± 1.39) vs Null (33.00 ± 2.34), p=0.011. (F, G) Nerve cross sections from Gli1 het and null mice injected intravenously with Evans Blue Albumin (EBA, red) shows the dye is contained within endoneurial vessels in both Gli1 hets and nulls (arrowheads). Dye also diffuses along MFs in Gli1 nulls (arrow) but does not penetrate into the endoneurium. (H, I) Nerve sections from Gli1 het and null fate mapped mice were crossed with the CX3CR1^EGFP/+^ reporter line to label macrophages with GFP. (J) The density of macrophages was not significantly altered in Gli1 nulls (30.44 ± 3.25) vs hets (23.33 ± 2.99), p=0.18. Values shown represent mean ± SEM analyzed using unpaired t-test with Welch’s correction based on n=3 biological replicates per genotype; * p<.05, ** p<.01, ns=not significant. Scale bars: (A, B: main) 100 µm, (A, B: insets) 5 µm, (D) 50 µm, (F-I) 100 µm.

We next compared the ultrastructure of Gli1 het and null peripheral nerves by EM (Figures 9, 10) and by staining of semi-thin (1 µm) sections with Toluidine Blue (data not shown). A striking finding is a strong reduction in fibrillar collagen in the ECM of the Gli1 null endoneurium (Figure 9A-C). In agreement, staining for collagen with either Masson’s Trichrome (Figure 9D, E) or with an antibody against Collagen-1 (Figure 9F, G) was substantially reduced in the Gli1 nulls. EFs largely generate non-basal lamina-associated, fibrillar collagens via their expression of the salient collagen transcripts (Col1a1, Col1a2, Col3a1) and high levels of prolyl-4-hydroxylase (Richard et al., 2014; Carr et al., 2019), an enzyme critical for collagen synthesis. We confirmed there is a significant decrease in Col1A1, 1A2, and 3A1 transcripts in Gli1 null nerves by qPCR (Figure 9H). Thus, a decrease in EF collagen synthesis accompanies their phenotypic shift towards MF formation.

**Figure 9:**
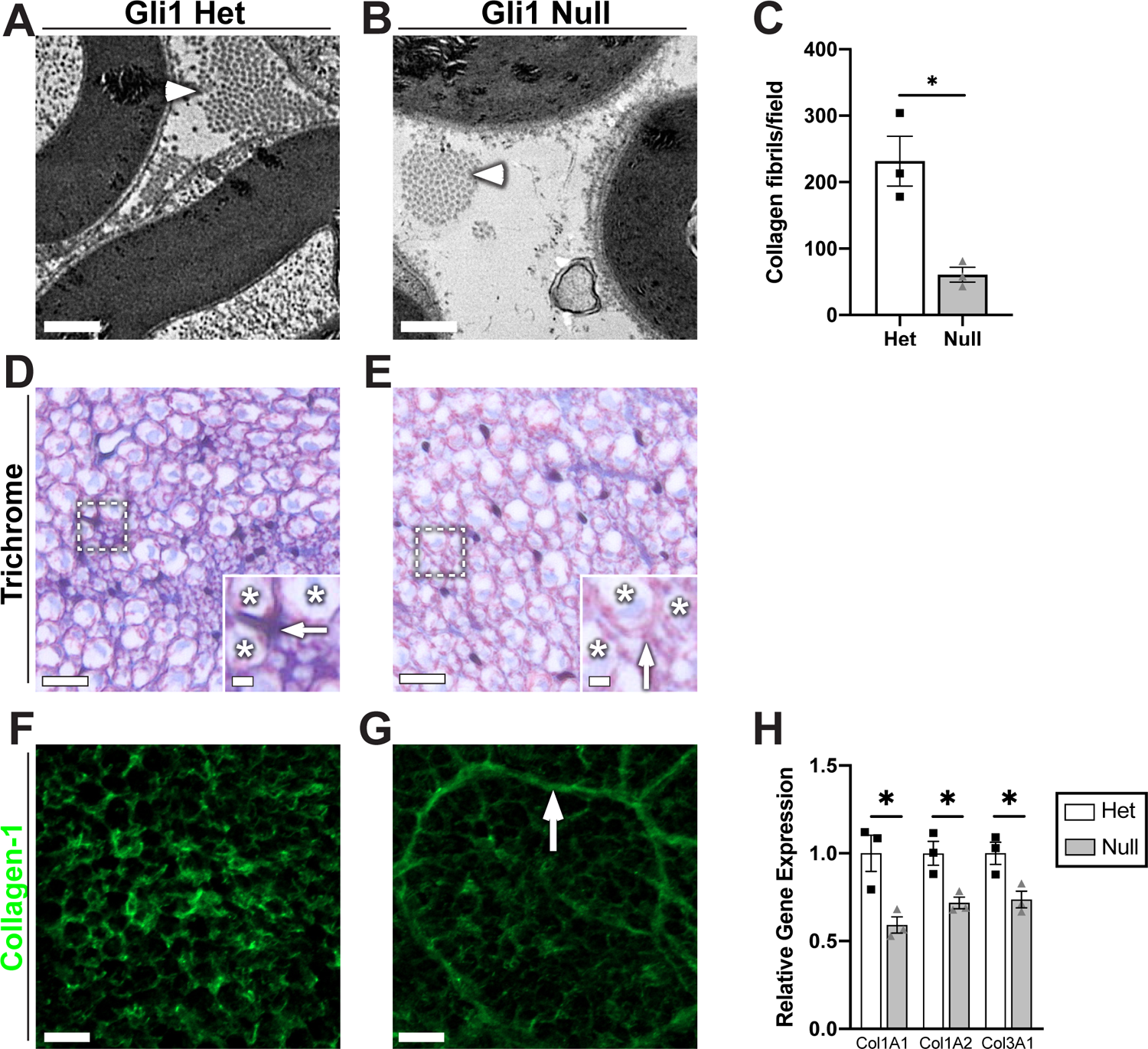
Gli1 expression controls peripheral nerve extracellular matrix (A, B) High magnification EMs reveal a decrease in collagen fibrils (arrowheads) in the Gli1 nulls vs. hets. (C) Quantification of collagen fibril density demonstrates a significant decrease in Gli1 nulls (60.67 ± 11.05) compared to hets (231.7 ± 37.55), p=0.036. (D, E) Gli1 het and null sciatic nerves were stained with Masson’s Trichrome, which labels collagen blue. Staining is decreased in the extracellular spaces (insets, arrows) between axons (insets, stars) of Gli1 nulls. (F, G) Gli1 het and null nerves were stained with an antibody against full-length (native) collagen-1 (green). Staining intensity is decreased in the endoneurium of Gli1 nulls although MF are moderately stained (arrow). (H) Transcript levels for several key collagen genes were determined by qPCR and show significant decreases for the 3 main fibril-forming collagen genes expressed in the endoneurium. COL1A1: Het (1.00 ± 0.10) vs Null (0.59 ± 0.05), p=0.042; COL2A1: Het (1.00 ± 0.07) vs Null (0.72 ± 0.033), p=0.035; COL3A1: Het (1.00 ± 0.06) vs Null (0.74 ± 0.05), p=0.033. Values shown represent mean ± SEM analyzed using unpaired t-test with Welch’s correction based on n=10 fields per replicate (C) and n=3 biological replicates per genotype (C, H); * p<.05. Scale bars: (A, B) 0.5 µm, (D, E, main) 10 µm, (D, E, insets) 2 µm, (F-G) 10 µm.

**Figure 10:**
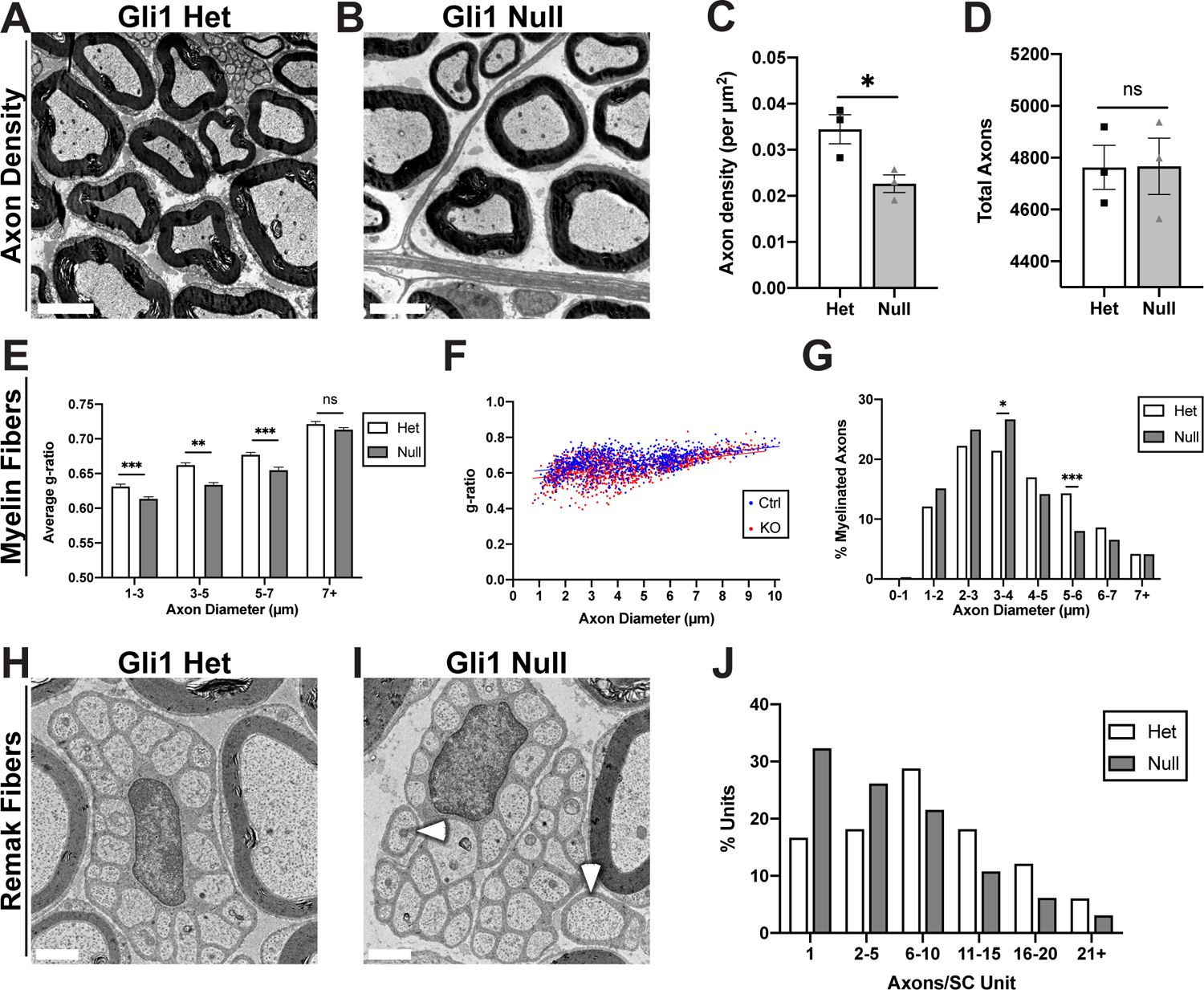
Gli1 expression regulates the morphometry of axon-Schwann cell units (A, B) EM cross sections demonstrate increased spacing between axon-SC units in nerves of Gli1 nulls vs hets. (C) Calculation of axon density reveals a significant decrease in Gli1 nulls (0.023 ± 0.0019) compared to hets (0.034 ± 0.0031), p=0.0424. (D) Total axon numbers were similar in Gli1 het (4763 ± 85.38) and null nerves (4564 ± 108.9), p=0.98. (E) Average g-ratios of myelinated fibers were calculated within the indicated diameter ranges and indicate a modest hypermyelination of small and medium size axons in Gli1 nulls. 1-3 µm: Het (0.63 ± 0.004) vs Null (0.61 ± 0.003), p=0.003, 3-5 µm: Het (0.66 ± 0.003) vs Null (0.63 ± 0.003), p<.0001, 5-7 µm: Het (0.67 ± 0.004) vs Null (0.65 ± 0.004), p=0.0001, 7+ µm: Het (0.72 ± 0.003) vs Null (0.71 ± 0.003), p=0.081. (F) Scatterplot of axon diameter vs g-ratio for single myelinated fibers in adult Gli1 hets and nulls. Distributions are largely overlapping with a slight shift towards lower g-ratios for smaller-diameter axons for Gli1 nulls vs hets. Lines of best fit: Gli1 het (y=0.015x + 0.60) vs Gli1 null (y=0.017x + 0.56). (G) A shift from larger to smaller axon diameters is observed in Gli1 nulls. p=0.63 (Total vs 0-1 µm), p=0.12 (Total vs 1-2 µm), p=0.31 (Total vs 2-3 µm), p=0.042 (Total vs 3-4 µm), p=0.18 (Total vs 4-5 µm), p=0.0002 (Total vs 5-6 µm), p=0.13 (Total vs 6-7 µm), p=0.99 (Total vs 7+ µm). (H, I) Remak bundles containing as few as 1-2 axons (I, arrowheads) were frequently observed in Gli1 nulls but not in hets. (J) Remak bundles were binned by number of axons per bundle, showing a shift towards lower axon:SC ratios (i.e., 1:1 and 1:2) in Gli1 nulls. (C, D) Values shown represent mean ± SEM analyzed using unpaired t-test with Welch’s correction based on n=3 biological replicates per genotype; *p<.05, ns=not significant. (E-G) >1000 axons across n=3 biological replicates per genotype were traced, (E) Axons were binned by diameter and average g-ratio was analyzed by t-test as above, (G) Axon diameters were binned as a percentage of total axons measured and analyzed by Fisher’s exact test; * p<.05, ** p<.01, *** p<.001. (J) >150 Remak bundles per genotype were analyzed, number of observations was insufficient for Fisher’s testing. Scale bars: (A, B) 5 µm, (H, I) 1 µm.

### Gli1 regulates axon-Schwann cell units non-autonomously

Along with a reduction in the ECM, there were significant changes in axon-SC units. These included a significant decrease in the packing density of axon-SC units in Gli1 null nerves compared to hets as evident by EM (Figure 10A-C). This decrease in density in the nulls reflects increased spacing, not axonal loss as the numbers of axons in the het and null nerves were equivalent (Figure 10D). In agreement, the nerves in the Gli1 nulls were nearly 50% larger in their cross-sectional area (data not shown).

In addition to altered density, there were also modest changes in the ultrastructure of axon-SC units. Myelin sheaths were, on average, slightly thicker in the Gli1 nulls indicated by a decrease in the g-ratios for small- and medium-diameter myelinated axons in the Gli1 nulls compared to hets (Figure 10E, F). There was also a modest (leftward) shift in the distribution of axon diameters in Gli1 nulls with increased numbers of smaller and fewer numbers of large-diameter axons (Figure 8G). Finally, there is a bias towards lower SC:axon ratios (i.e., more 1:1 and 1:2 vs. fewer 1:11+) in Remak fibers of Gli1 nulls than hets (Figure 10H-J). Given that neither neurons nor SCs express Gli1, these latter findings suggest a non-autonomous role of Gli1, likely via EFs, in regulating the precise ultrastructure of axon-SC units.

### Gli1 maintains the endoneurial organization of adult peripheral nerves

These studies indicate that Gli1 regulates the organization of the endoneurium during development. An important question is whether Gli1 is also required to maintain this organization in adult nerves. To address this question, we first generated a floxed allele of Gli1 as shown schematically in Figure 11A. The fidelity of the insertion of loxP sites was corroborated by sequencing. Mice with the floxed Gli1 allele were crossed to the *Gli1^CE/+^* driver and the Ai9 reporter to generate mice with *Gli1^CE/Fl^;Ai9* and *Gli1^CE/+^;Ai9* genotypes, enabling Gli1 fate mapping of inducible KOs (iKOs) and hets, respectively. Specifically, mice were treated with tamoxifen on alternate days (x4) and analyzed 5 and 8 weeks later (Figure 11B). In the absence of tamoxifen, there was no reduction of Gli1 levels in the peripheral nerves of *Gli1^CE/Fl^;Ai9* mice based on qPCR. In contrast, 5 and 8 weeks after tamoxifen treatment, Gli1 levels in iKO sciatic nerves were reduced to 1-2% of littermate controls (Figure 11C).

**Figure 11:**
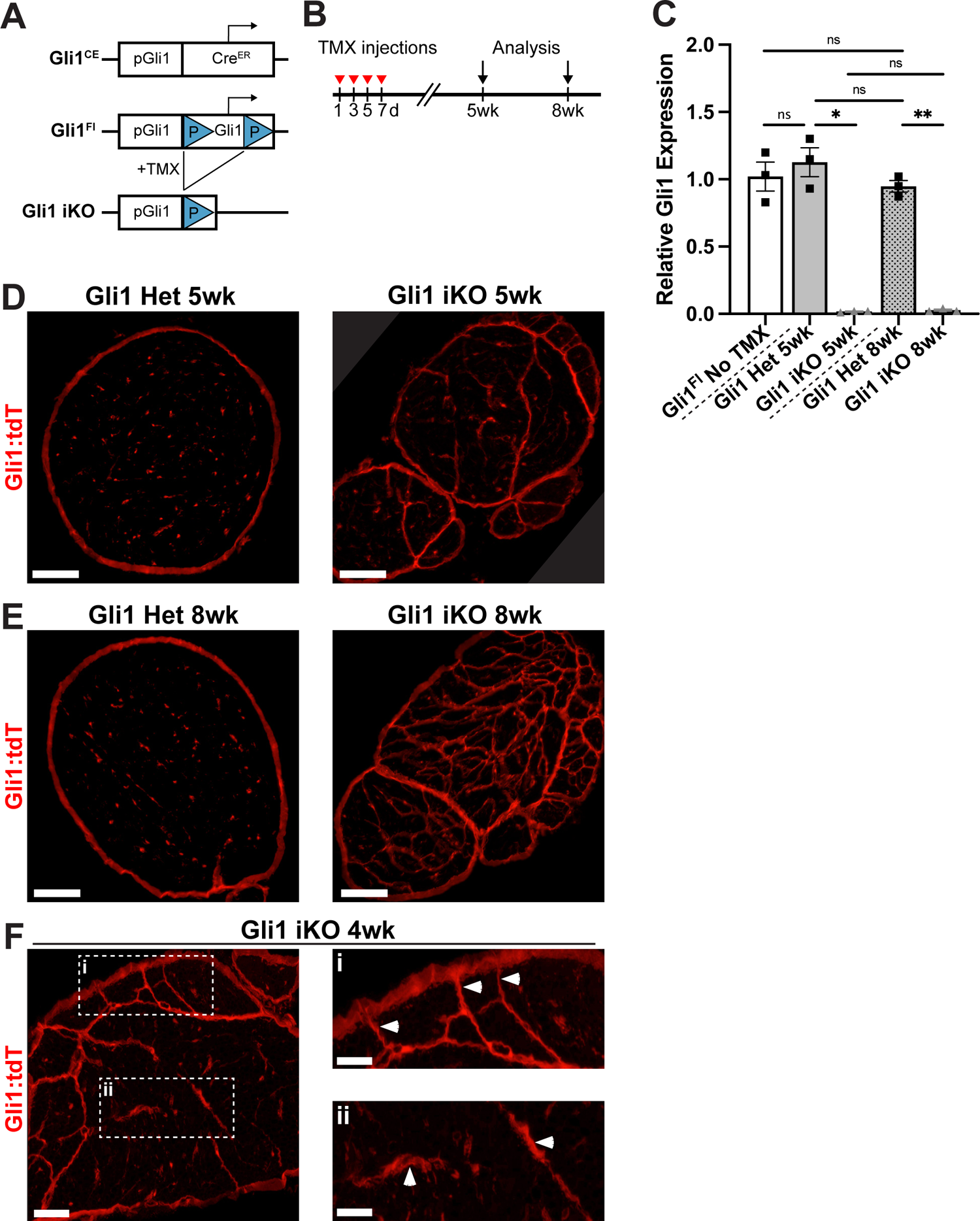
Inducible knockout of Gli1 results in minifascicle formation in adult nerves (A) Schematic of Gli1^CE^ and Gli1^Fl^ alleles. LoxP sites flank exons 5-9 of the Gli1 coding sequence. After tamoxifen administration to Gli1^CE/Fl^ mice, Cre-mediated excision of this sequence results in Gli1 inducible knockout (iKO). Gli1^CE^ also drives expression of the Ai9 reporter. (B) Tamoxifen was given on alternating days for 4 injections and analyses were performed at 5 and 8 weeks. (C) qPCR for Gli1 transcripts reveals equivalent levels in Gli1^CE/Fl^ mice prior to tamoxifen administration and Gli1^CE/+^ hets at 5 and 8 weeks, confirming that insertion of the LoxP sites did not interfere with endogenous Gli1 transcription. In Gli1 iKO mice at both 5 and 8 weeks, Gli1 transcript levels are reduced to 1-2% of littermate controls. Gli1^CE/Fl^ No TMX (1.00 ± 0.11), Gli1 Het 5 wk (1.13 ± 0.11), Gli1 iKO 5 wk (0.017 ± 0.003), Gli1 Het 8 wk (0.95 ± 0.04), Gli1 iKO 8 wk (0.027 ± 0.007); Het 5 wk vs iKO 5 wk p=.03, Het 8 wk vs iKO 8 wk p=.008. (D) Gli1 het and iKO nerves at 5 weeks post-tamoxifen shows fate mapped cells forming nascent MFs. (E) By 8 weeks post-tamoxifen, Gli1 iKO nerves contain dense MFs, similar to Gli1 constitutive nulls. (F) Examination at an early timepoint (4 weeks) shows MFs forming both at the periphery (inset i) as well as in the interior (inset ii) of the nerve. Values shown in panel C represent mean ± SEM analyzed using Welch’s ANOVA test with Dunnett’s T3 multiple comparisons test based on n=3 biological replicates per genotype; * p<.05, ** p<.01, ns=not significant. Scale bars: (D-E) 100 µm, (F, main) 50 µm, (F, insets) 25 µm.

Nerves from Gli1 iKOs revealed a striking reorganization of the nerve by MFs, which were evident at 5 weeks and robust at 8 weeks (Figure 11D and E, respectively). There were no evident changes in the perineurium of Gli1 iKOs based on fate mapping. At 3.5 to 4 weeks, the earliest times examined, MFs were still forming (Figure 11F). Many of the MFs present at these earlier times were locally peripherally and connected to the perineurium (Figure 11F, inset i). However, there were also multiple examples of forming MFs that appeared to be present entirely within the endoneurium that lacked an evident connection to the perineurium (Figure 11F, inset ii). This latter observation suggests existing EFs within the endoneurium can reorganize to drive MF formation. In complementary studies, we investigated whether re-expression of Gli1 would revert MFs that had formed in the constitutive Gli1 nulls. We crossed *Gli1^CE^* mice to *Rosa26^flox-stop-flox-FLAG-Gli1^* (*Rosa^Gli1^*) mice (Vokes et al., 2007). These mice were further crossed to a GFP reporter i.e., *Rosa26^CAG-EGFP^* (RCE) allele (Figure 12A). Tamoxifen administration on alternate days (x4) enabled GFP fate mapping of Gli1+ cells in hets and nulls while simultaneously driving expression of a FLAG-tagged Gli1 allele in the same cells. Analysis was performed 6 weeks later (Figure 12B). Based on qPCR, Gli1 expression was restored in Gli1 null mice that overexpressed the FLAG-Gli1 allele (Gli1 null OE) to 91% of the levels of hets (Figure 12C). There was also a dose-dependent, increase in Gli1 expression in Gli1 het OE mice, i.e., 81% greater than in het controls. Overexpressing Gli1 in hets had no overt effect on the organization of the nerves (Figure 12D). Of note, despite restoration of Gli1 to essentially normal levels in the Gli1 null OE nerves, there was no reduction in MFs or morphological changes in the Gli1-expressing cells compared to Gli1 nulls (Figure 12E), even out to 10 weeks, the latest time point examined (data not shown).

**Figure 12:**
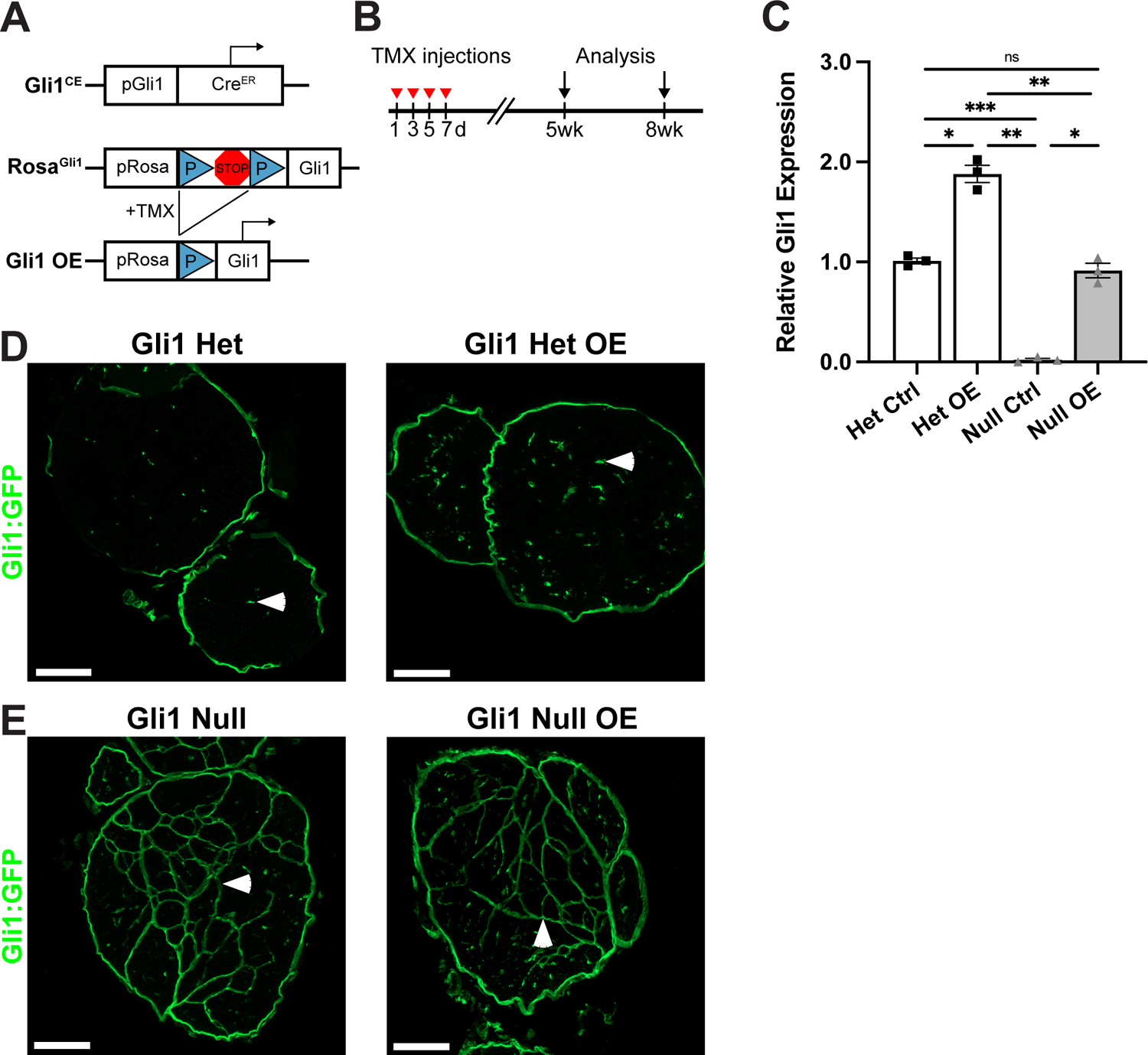
Restoring Gli1 expression in Gli1 nulls does not revert minifascicles (A) Diagram of Gli1^CE^ and ROSA^Gli1^ alleles. LoxP sites flank a transcriptional stop site upstream of the full Gli1 coding sequence. After tamoxifen administration to Gli1^CE^; ROSA^Gli1^ mice, Cre-mediated excision of the stop site results in Gli1 overexpressors (OE) which constitutively express Gli1 under the ROSA26 promoter. Gli1^CE^ also drives expression of the RCE (GFP) reporter. (B) Tamoxifen was given on alternating days for 4 injections and analysis was performed at 6 weeks. (C) qPCR was performed on sciatic nerves of adult Gli1 het ctrl (*Gli1^CE/+^;RCE),* null ctrl (*Gli1^CE/CE^;RCE)*, het OE (*Gli1^CE/+^;RCE/RGli1),* and null OE (*Gli1^CE/CE^;RCE/RGli1)* mice. Gli1 transcript levels in Het OE are increased 81% above Gli1 het controls. Gli1 null controls express virtually no Gli1, while Gli1 null OE have transcript levels 91% of het controls. Het Ctrl (1.00 ± 0.029), Het OE (1.88 ± 0.087), Null Ctrl (0.022 ± 0.014), Null OE (0.91 ± 0.072); Het Ctrl vs Null Ctrl p=.0003, Het Ctrl vs Het OE p=0.033, Null ctrl vs Null OE p=0.02, Null ctrl vs Het OE p=0.0068. (D-E) GFP staining of fate mapped cells in the indicated genotypes shows no evident morphological differences between het ctrl and het OE (D) or between null ctrl and null OE (E). Isolated EF are labeled in all Gli1 hets (D, arrowheads) and MFs are labeled in Gli1 nulls (E, arrowheads). Values shown in panel C represent mean ± SEM analyzed using Welch’s ANOVA test with Dunnett’s T3 multiple comparisons test based on n=3 biological replicates per genotype; * p<.05, ** p<.01, *** p<.001, ns=not significant. Scale bars: (B-E) 100 µm.

Taken together, these results indicate that loss of Gli1 is sufficient to drive formation of MFs in healthy adult nerves over several weeks. In contrast, MFs formed during development are quite stable even when Gli1 levels are restored to essentially normal levels during the time interval examined.

## DISCUSSION

We have shown that EFs, PNG, and pericytes are Gli1-expressing cells. While loss of Gli1 did not affect development or maintenance of the perineurium, it strongly impacted the endoneurium, notably resulting in formation of MFs during development and in adult nerves following iKO. These MFs are similar to those previously described in Dhh nulls (Parmantier et al., 1999), a similarity that initially suggested Gli1 as a candidate effector of Dhh in the endoneurium. Unexpectedly, Gli1 is persistently expressed in the EFs of Dhh nulls. As Dhh is the principal hedgehog in the PNS, these results indicate non-canonical pathways drive expression of Gli1 in EFs. The presence of MFs in both Gli1 and Dhh nulls despite ongoing Gli1 expression in the latter also indicates that Gli1 and Dhh function non-redundantly. Gli1 must therefore act in concert with other downstream effectors of Dhh to drive acquisition of normal endoneurial architecture and preclude EFs from organizing into MFs. In contrast, formation of the perineurium does not require Gli1 but rather must be mediated via other effectors of Dhh, e.g., via canonical signaling. Our results also extend the roster of reciprocal signaling between different cells types in peripheral nerves. Thus, Gli1 null nerves exhibit altered ECM composition, likely due to dysregulation of EFs, as well as non-autonomous effects on vascular organization and the morphometry of axon-SC units. We consider these points further below.

### Gli1 is expressed by cells with diverse embryologic origins

Here, we have identified EFs, PNG, and pericytes as distinct Gli1-expressing cells in peripheral nerves, extending a previous study that identified EFs as Gli1-expressing cells (Bobarnac Dogaru et al., 2018). These three cell types have distinct embryological origins: EFs arise from neural crest stem cells (Joseph et al., 2004) whereas PNG derive from Nkx2.2-positive precursors that egress from the ventral spinal cord (Kucenas et al., 2008; Clark et al., 2014). The origins of peripheral nerve pericytes remain to be established.

Gli1-positive precursors of these various populations are present in developing peripheral nerves as early as E12-E15 based on *in utero* fate mapping (Figure 4A, B), consistent with EM analysis of the cellular composition of the developing PNS (Jessen and Mirsky, 2005). SCs were not labeled by the *in utero* fate mapping indicating that early neural crest stem cells and their glial-restricted progeny, which express Dhh starting at E12 (Jaegle et al., 2003), are not subject to autocrine Hh signaling. These results, together with a prior study (Joseph et al., 2004), underscore that distinct neural crest-derived precursors give rise to SCs and Gli1-positive EFs and that these lineages become restricted prior to E15. In addition to the SC and EF lineages, a Gli1-positive precursor that is not neural crest-derived is also present at this time and gives rise to PNG and/or pericytes. Given that the overall nerve architecture of Gli1 hets and nulls is similar at P1 but diverges significantly by P10 (Figure 4C-F), Gli1 expression is dispensable for the generation of these various precursor populations *in utero* and functions principally in the early postnatal period. Despite their distinct embryologic origins, EFs and PNG express a number of molecular markers in common in addition to Gli1. A recent single-cell RNA-sequencing analysis of PNS cells found PDGFR*α* to be a broad marker of non-SC populations in the PNS - principally EFs and PNG (Carr et al., 2019). Transcriptionally unique populations of EFs, PNG, and epineurial fibroblasts were identified by this analysis. Querying this data set reveals that a significant portion of each of cell type is indeed Gli1-positive. These results are therefore consistent with the nearly uniform co-labeling of cells by Gli1 and PDGFR*α* (Figure 1D). In the future, fate mapping using markers unique for each of these populations will enable better delineation of their embryologic origins and lineage relationships than is currently possible.

### Non-canonical expression of Gli1 cooperates with Dhh signaling to regulate normal endoneurial development

As noted, we initially considered Dhh released by mSCs to be a likely driver of Gli1 expression in PNG and EFs. First, loss of Egr2 results in an overall decrease in Dhh with a commensurate decrease in Gli1 transcripts by qPCR (Figure 2E). Second, loss of Gli1 phenocopies much of the reported endoneurial phenotype of Dhh nulls: e.g., reorganization into MFs (Figure 3A, B), increased capillary density (Figure 8A-E), and defective ECM (Figure 9) (Parmantier et al., 1999; Chapouly et al., 2016). Third, PNG and EFs express the Patched-1 receptor (Parmantier et al., 1999; Sharghi-Namini et al., 2006) and Gli2 (Extended Data Figure 1) - the latter is considered an obligate canonical activator of Gli1 expression. Finally, EFs treated with Shh ligands *in vitro* upregulate Gli1 expression (Bobarnac Dogaru et al., 2018).

However, our data indicates expression of Gli1 in EFs is unlikely to be downstream of Dhh given that Gli1:*β*Gal continues to be expressed in the EFs of Dhh nulls (Figure 2F, I). Gli1 transcripts persist at 44% of control levels in P5 Dhh KO nerves, which lack detectable expression of any Hh family members, strongly supporting Hh-independent expression of Gli1 in these cells (Figure 2H). Determining the extent of the reduction of Gli1 expression in the P5 Dhh null EFs, or if there is any reduction at all, is confounded in this qPCR analysis by the near-absence of PNG and epineurial cells. These latter two cell populations likely contribute substantially to the Gli1 levels measured in control nerve lysates suggesting that the loss of Gli1 in EFs in the P5 Dhh nulls may be nominal. Indeed, starting by at least P25, Dhh nulls exhibit normal or even increased levels of Gli1 (Figure 2K) further underscoring that Gli1 expression is independent of Dhh. A modest increase may result from increased numbers of Gli1-positive cells akin to that seen in the Gli1 nulls (Fig. 5C) We also considered neurons as a potential source of hedgehog ligands within peripheral nerves. Shh is expressed by a subset of dorsal root ganglion neurons and is transported into their distal axons, driving Gli1 expression in the surrounding follicular epithelium (Brownell et al., 2011). This potential source of Shh would not be detected by qPCR of nerves as its transcripts/synthesis reside in the soma. However, Shh conveyed by axons is an unlikely driver of Gli1 in EFs in healthy nerves. First, it is not known if Shh can be released along the length of axons or if it is only released at their termini. Even if Shh is released *en passant*, axons are wrapped by Remak or mSC processes along their length, which would be expected to impede access to endoneurial cells. In follow-up studies, we carried out nerve transection to remove axons as a potential source of Hh ligands (Brownell et al., 2011) and 1 week later, Gli1:*β*Gal continued to be expressed in cells in the distal (data not shown). These latter results are potentially ambiguous as expression of Shh is reported to be upregulated in SCs following nerve injury (Hashimoto et al., 2008).

Thus, normal endoneurial development requires co-incident signaling from non-canonically expressed Gli1 and from other (non-Gli1) signals downstream of Dhh. The ligands and pathways that drive non-canonical Gli1 expression in EFs of the Dhh nulls are not known. Candidate pathways include MAPK/ERK (Po et al., 2017; Hasan et al., 2020) and/or TGF*β*/SMAD signaling pathways (Dennler et al., 2007). The pathways downstream of Dhh likely include canonical, Smo-dependent generation of the Gli2A transcriptional activator or inactivation of the Gli3R repressor (Ingham and McMahon, 2001). Gli1 and Gli2 were previously shown to have significant functional redundancy *in vivo* (Bai and Joyner, 2001) and Gli2 is known to physically and functionally interact with Gli1 to coordinately regulate transcription (Tolosa et al., 2020), suggesting a mechanism for coincident canonical and non-canonical control of EF differentiation, respectively.

### Formation and Stability of Minifascicles

A long-standing question is what is the cell of origin that gives rise to MFs. Prior studies had suggested PNG as a potential source of MFs based on the morphological and marker similarities of cells in MFs to PNG (Burkel, 1967; Parmantier et al., 1999). Results from this study indicate that EFs are a much more likely cellular source of these structures. In the Dhh nulls, there is almost no perineurium (Figure 2F), yet despite the absence of PNG, formation of MFs is robust (Figure 2I). In Gli1 nulls, the perineurium appears normal (Figure 3E), yet endoneurial not perineurial, Gli1-positive cells exhibit increased proliferation in these mice (Figure 5D). There are also decreased numbers of free EFs in the Gli1 nulls (Figure 5G). These latter results suggest dividing EFs are recruited to form MFs. In addition, EFs undergo a morphological transformation postnatally in the Gli1 nulls. Unlike Gli1 het EFs, which gradually retract processes potentially in response to Dhh released postnatally by mSCs, these processes persist and extend further in the Gli1 nulls, eventually interdigitating with each other (Figure 4F, H). A similar morphological transformation of EFs was evident in adult Gli1 iKOs at the onset of MF formation, where they often lacked an apparent connection to the perineurium (Figure 11F). Compellingly, MFs that initially form in both the Dhh and Gli1 nulls express EF but not PNG markers (Figure 7).

Taken together, these results strongly support an EF origin of MFs. As MFs become more robust over time, they express PNG markers (e.g., Glut-1), and become fully integrated with the perineurium. Expression of PNG markers in mature MFs could reflect invasion of MFs by PNG over time. Alternatively, EFs may upregulate PNG markers as MFs mature due to density-dependent effects on expression (Extended data Figure 2). Consistent with the latter possibility, i.e., that EFs can acquire PNG characteristics, EFs appear able to regenerate a full perineurial sheath following microsurgical stripping of the perineurium (Nesbitt and Acland, 1980). Similarly, purified fibroblasts added to myelinating SC/neuron cocultures form perineurial-like structures (Bunge et al., 1989).

Following injury/transection, MFs surround SC basal lamina tubes (i.e., Bungner Bands) in the distal stumps (Morris et al., 1972; Röyttä et al., 1987). Our findings thus suggest resident EFs are likely to form MFs after nerve injury (Röyttä et al., 1987; Röyttä and Salonen, 1988; Vallat et al., 1988) or in neuropathies (Baldinotti et al., 2018). An EF source is consistent with a prior study showing that MFs assemble in transected rat sciatic nerves despite prior mechanical removal of the local perineurium (Terho et al., 2002). Recent data demonstrates MFs in injured nerves arise from Gli1-positive cells (Bobarnac Dogaru et al., 2018; BZ, JLS unpublished). Future studies using cell specific markers will be useful to determine if EFs also form injury-induced MFs.

We have also shown that iKO of Gli1 in adult nerves is sufficient to drive MF formation within several weeks (Figure 11). This result supports the notion that reductions of Dhh and/or Gli1 signaling that ensue with injury/SC dedifferentiation contribute to MF formation in pathological settings. Once formed, MFs appear quite stable as re-expression of Gli1 in adult Gli1 nulls failed to reverse the MF phenotype after 10 weeks (Figure 12). Previous work has shown that MFs formed following nerve transection injury are also very stable. MFs tend to disappear ∼20 weeks with successful reinnervation of the transected nerve (Röyttä and Salonen, 1988), suggesting that these structures can undergo slow disassembly with SC redifferentiation driven by axonal signals.

### Potential effects of MFs on peripheral nerve function

The findings in the Gli1 nulls raise the question of whether there are pathological or functional consequences of reorganizing nerves into MFs. Normally, motor and sensory fibers in peripheral nerves maintain a high degree of somatotopy despite frequent anastomoses between perineurial sheaths. Thus, motor fibers for specific muscle groups and sensory fibers for specific cutaneous areas tend to remain grouped as fascicles or within fascicles along the length of peripheral nerves (Badia et al., 2010; Mioton et al., 2019). In the case of the Gli1 nulls, SBF-SEM revealed that the cellular sheaths forming MFs are not confined to a fixed set of axons but rather anastomose along their length (see Extended Data Video 1). As the Gli1 nulls do not exhibit any gross functional deficits that would be expected from mistargeting of axons, fascicles seem likely to retain their normal somatotopy despite subdivision into fascicles that contain as few as 1-2 axons.

Prospective functional consequences of MFs have been considered primarily in the context of nerve injury and are extrapolated from the normal functions of the perineurium, which they resemble. MFs are thought to restore the tissue barrier function that is lost with the disruption of the perineurium in mechanical injury (Ahmed and Weller, 1979). In agreement, MFs form tight junctions and provide a competent barrier against infiltration of injected tracers into fascicles (Figure 3G-I). Akin to the perineurium, MFs are likely to protect against stretch injury (Schraut et al., 2016), which damaged peripheral nerve/distal stumps may be particularly susceptible to. Also similar to the perineurium, MFs may protect nerve fibers from immunologic damage, including complement, and help maintain homeostatic intraneural pressure and metabolism (Zochodne, 2009; Weerasuriya and Mizisin, 2010).

### Nerve sheath and barrier development is Dhh-dependent but Gli1-independent: implications for axonal degeneration

A striking finding of this study is that while Dhh nulls exhibit major defects of the perineurium and epineurium (Parmantier et al., 1999), the Gli1 nulls do not (Figure 3D-F). Thus, PNG development requires Dhh signals potentially via canonical signaling that is Gli1-independent, e.g., via Gli2A. In agreement, a recent study examining regeneration after nerve injury with a *Gli1^CreERT2^* driven cKO of Smo reported the perineurium failed to reform (Yamada et al., 2021). Whether expression of Gli1 itself is canonical or non-canonical in PNG is unclear as there are essentially no PNG to monitor in the *Dhh^-/-^; Gli1^LacZ/+^*mice. Future studies with inducible KOs of Smo or Dhh after the perineurium has formed will be useful to determine if Gli1 expression in PNG is canonical or not.

An additional difference between Dhh and Gli1 nulls is that the former exhibit axonal degeneration/regeneration with increased age (Sharghi-Namini et al., 2006) whereas Gli1 nulls do not, even at 18 months (data not shown). This difference suggests the aberrant perineurium of the Dhh nulls, with its associated loss of BNB function (Sharghi-Namini et al., 2006), underlies the axonal degeneration in these mice rather than the re-organization into MFs or ECM changes, which are seen in both Dhh and Gli1 nulls. Likewise a defective perineurium and BNB may account for the axonal injury and severe peripheral neuropathy seen in patients with Dhh mutations, despite their designation as minifascicle neuropathies (Baldinotti et al., 2018). Macrophage numbers are increased in the Dhh (Sharghi-Namini et al., 2006) but not significantly in the Gli1 nulls (Figure 8H-J) nerves, potentially resulting from the aberrant BNB in the former. Whether this inflammation contributes to and/or is a consequence of axonal degeneration in these animals is not yet known.

### Gli1 demarcates peripheral nerve pericytes and regulates vascular organization

There is a striking expansion of the nerve vasculature in Gli1 nulls which is closely associated with MF formation. The perivascular Gli1+ cells in the PNS are plausible candidates to direct this remodeling of the vasculature as they are known to directly regulate angiogenesis by developing ECs (Bergers and Song, 2005). Indeed, in a recent study (Chen et al., 2020), genetic ablation of Gli1-positive perivascular mesenchymal stem cells surrounding capillaries in bone resulted in defective angiogenesis in both healthy and injured bones. This pro-angiogenic effect is believed to be mediated by HIF-1*α* signaling (Chen et al., 2020). Additionally, a recent report identified Gli1-expressing EFs as a source of VEGF in the setting of nerve injury (Faniku et al., 2021). Our data are consistent with this possibility given the close association of the EF-derived MFs and the vasculature (Figure 8A-E and Extended data video 1). The role of this increased vascularization is not known, but may provide enhanced metabolic support to nerves partitioned by MFs by compensating for any constraints on nutrient diffusion within the endoneurium imposed by MFs. In potential agreement, increasing angiogenesis via VEGF gene therapy improves axonal survival following injury (Pereira Lopes et al., 2011).

### Myelinating SCs regulate Gli1 levels and assembly of peripheral nerve compartments

We have corroborated that Egr2 activity regulates Dhh production by adult SCs, in agreement with a prior report (Jang et al., 2006). Thus, Dhh levels are reduced by ∼50% in Egr2 conditional KOs (Fig. 2E). This residual expression in the Egr2 nulls indicates other transcriptional activators must also regulate Dhh. One such transcription factor is Sox10, which directly regulates Dhh expression (Küspert et al., 2012), and thereby the perineurium, and cooperates with Egr2 to drive myelination.

As noted, there is also a commensurate reduction of Dhh and Gli1 in the Egr2 nulls. These results indicate Gli1 expression in the perineurium and endoneurium is regulated by factor(s) released from mSCs, coordinating Dhh and Gli1 expression with myelination. Interestingly, in the adult Egr2 nulls, EFs partially but incompletely organize as MFs (Figure 2D). This incomplete MF formation is consistent with significant, residual expression of both Dhh and Gli1 in these mice potentially driven by Sox10 expression. In contrast, MFs are a prominent feature of many dysmyelinating SC mouse mutations (Feltri et al., 2002; Darbas et al., 2004; Yu et al., 2005; Grove et al., 2007; Monk et al., 2011). Substantial reductions of Egr2 in these mouse mutants, with resulting reductions of Dhh and Gli1, likely contributes to their formation of MFs.

### Reciprocal intercellular signaling during PNS development

Our results, together with prior reports, underscore the rich, reciprocal set of intercellular signals that coordinates development of the various cellular compartments of peripheral nerves. Thus, axonal signals, including neuregulin, upregulate Egr2, which cooperates with Sox10 to initiate myelination (Kao et al., 2009; He et al., 2010). This in turn drives SC production of Dhh and upregulation of Gli1 in cellular targets to regulate nerve fasciculation, development of the vasculature, and ECM production. Our results further indicate these endoneurial changes reciprocally feedback to regulate axon-SC units. This is evidenced by the smaller axons, thicker myelin, and enhanced segregation of axons in Remak fibers in the Gli1 nulls (Figure 10).

Similar effects on myelinated axon diameters and Remak sorting, but not myelination, were described previously in Dhh mutants (Sharghi-Namini et al., 2006). The changes in Remak bundles in the Dhh nulls were considered a direct effect of Dhh on Remak SCs signaling via Ptch2 (Bajestan et al., 2006). As neither mSCs or Remak SCs express Gli1, any effects on myelinated and Remak fibers in the Gli1 nulls - and by inference potentially in the Dhh mutants - must be SC non-autonomous. Thus, signals originating in Gli1 null EFs, pericytes, and/or PNG must account for the altered morphology of these axon-SC units. This notion agrees with earlier studies in zebrafish that implicate PNG in regulating early SC development (Kucenas et al., 2008; Binari et al., 2013).

The signals from these other peripheral nerve components that regulate axon-SC units are unknown. ECM components released by EF and/or PNG are clearly reduced in the Gli1 nulls (Figure 9). These components may regulate nerve fiber packing density (Figure 10A-C) and contribute directly to the SC basal lamina or indirectly enhance its production (Obremski et al., 1993). The basal lamina is an important positive and negative regulator of SC myelination (Heller et al., 2014; Ghidinelli et al., 2017). Additionally, fibrillary collagens, which are reduced in Gli1 nulls, normally confer mechanical stability to peripheral nerves (Ushiki and Ide, 1986). Loss of these ECM components may alter nerve stiffness in the Gli1 nulls, impacting mechanical signaling that regulates SC differentiation (Sophie et al., 2019). Finally, there is a rich array of paracrine signals released by EFs, whose expression is affected by nerve injury (Toma et al., 2020), that may likewise be altered in the Gli1 nulls impacting SCs. A recent study identified EFs in regenerating nerves as a source of soluble neuregulin (Fornasari et al., 2020), an important signal that promotes the SC repair phenotype essential for nerve regeneration (Stassart et al., 2013). Future studies utilizing RNA-seq and proteomic analyses to elucidate further the nature of the peripheral nerve signals that impinge on SCs will be of considerable interest.

## ACKNOWLEDGEMENTS

The authors are grateful to the NYU Langone Health Microscopy Core for extensive assistance with electron microscopy, to Warren J. Leonard (NIH-NHLBI) for generously providing the *Egr2_flox_* mouse line, to the NYU Rodent Genetic Engineering Core for their assistance with generation of the *Gli1_flox_* mouse line, and to members of the Salzer lab for helpful discussions and comments on the manuscript. This work was supported by NS103353 to B.Z. and NS100867 to J.L.S.

## AUTHOR CONTRIBUTIONS

Conceptualization, B.Z. and J.L.S.; Methodology, B.Z., O.D., H.B., and J.S.; Investigation, B.Z., J.B., and O.D.; Writing – Original Draft, B.Z. and J.L.S.; Writing – Review & Editing, B.Z., H.B., J.S., and J.L.S.; Funding Acquisition, B.Z. and J.L.S.; Supervision, J.L.S.

---

**Extended Data Figure 1:**
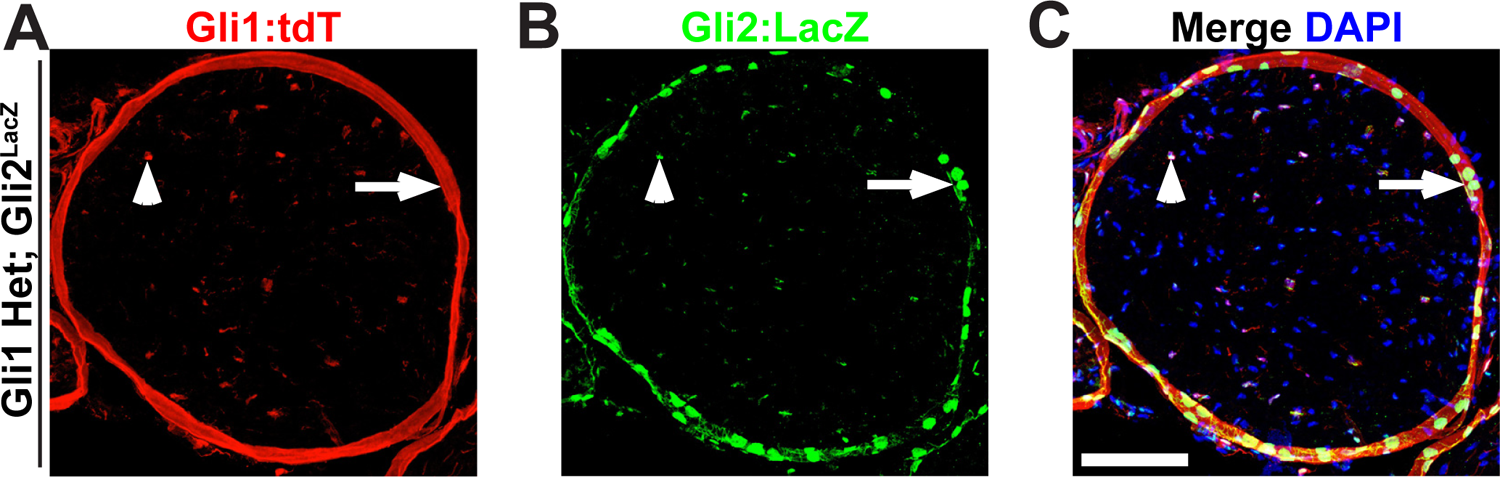
Gli2 is expressed in Gli1 fate mapped cells in the PNS (A, B) *Gli1^CE/+^;tdT* mice were crossed to *Gli2^nLacZ/+^* mice and fate mapped to label Gli1-expressing cells. Cross sections of sciatic nerves were stained for Gli1:tdT and *β*Gal (Gli2-positive cells, green). (C) The majority of endoneurial (arrowhead) and perineurial (arrow) Gli1 fate mapped cells co-expressed Gli2. Scale bars: (A-C) 100 µm.

**Extended Data Figure 2:**
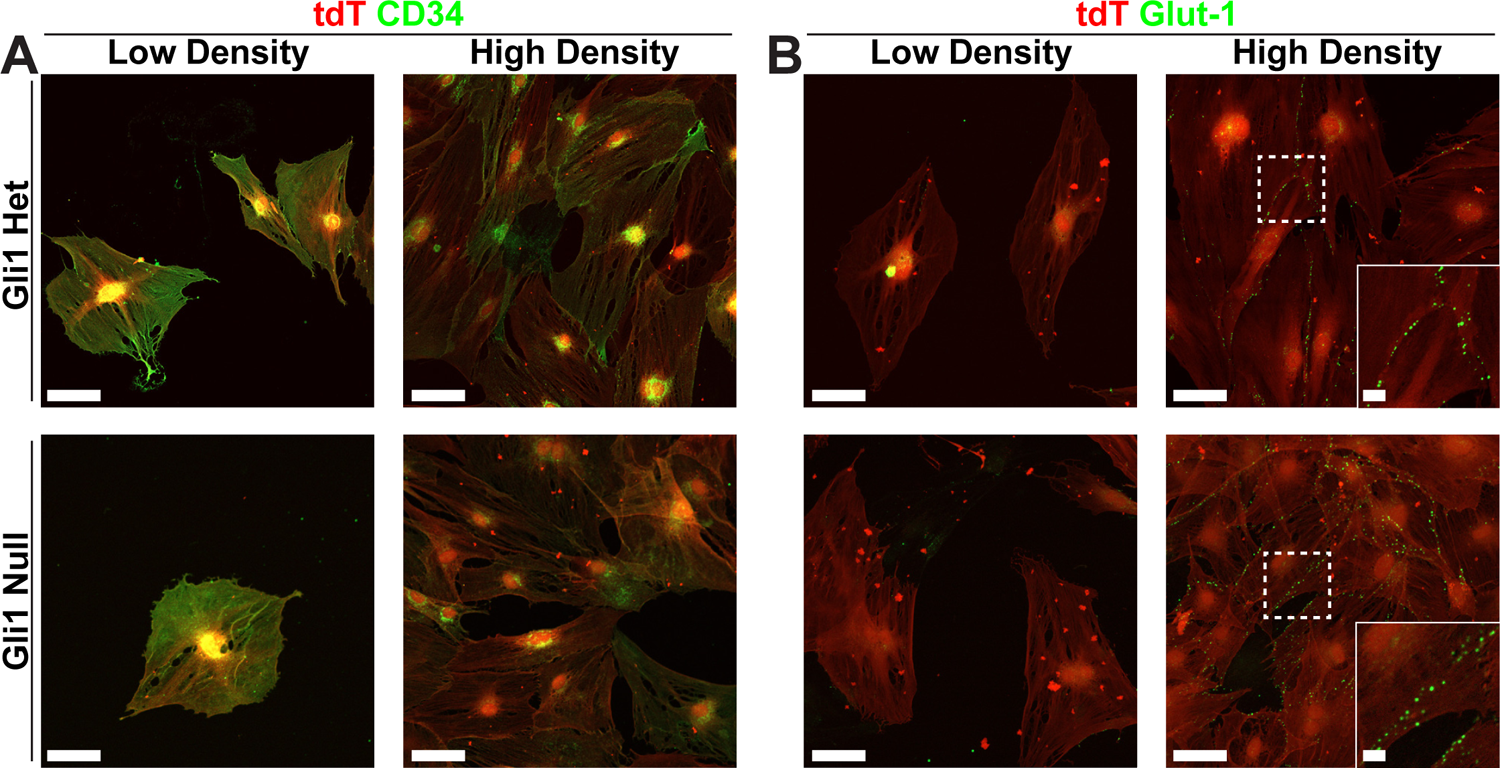
Expression of EF and PNG markers is regulated by cell density *in vitro* Fate mapped cells derived from sciatic nerve explants of Gli1 hets and nulls were plated at low density (2,000 cells/ cm^2^) or high density (10,000 cells/cm^2^) in standard EF media and fixed after 48 hr. (A) Cultures stained for Gli1:tdT (red) show robust expression of CD34 (green) in fate mapped cells at low density and an apparent decrease in expression when cells were grown at higher density. This effect was present in both Gli1 het and null cultures. (B) Cultures stained for Gli1:tdT (red) show no detectable staining of Glut-1 (green) in fate mapped cells grown at low density. At high density, however, punctate expression of Glut-1 is observed at sites of cell-cell contact in both het and null cultures (see insets). Scale bars (A-B, main) 50 µm, (B, insets) 10 µm.

## Extended Data

**Extended Data Video 1:** 3D view of an adult Gli1 null sciatic nerve (0:00-0:15) 3D reconstruction of adult Gli1 null sciatic nerves viewed in the XY plane and traversing along the z-axis. MF boundaries are traced in red, blood vessels (BV) are highlighted in blue. Frequent anastomoses of small MFs are seen with axons traversing between compartments. Note that BV are almost always embedded within MF structures. (0:15-0:30) 3D structures of MFs are viewed from the side. Next, axons/SC units are revealed. Finally, the BV network is highlighted and shown to run within MFs.

